# Neuropathological changes in the nucleus basalis of Meynert in people with type 1 or type 2 diabetes mellitus

**DOI:** 10.1101/2025.01.29.635423

**Authors:** Wei Jiang, Martin J. Kalsbeek, Felipe Correa-da-Silva, Andries Kalsbeek, Dick F. Swaab, Sarah E. Siegelaar, Chun-Xia Yi

**Author notes:** Correspondence: Chun-Xia Yi, MD PhD, Department of Endocrinology and Metabolism, Amsterdam University Medical Center, Location AMC, Meibergdreef 9, 1105 AZ, Amsterdam, E.

## Abstract

People with type 1 or type 2 diabetes mellitus (T1DM or T2DM) often experience cognitive impairment. We profiled cells in the nucleus basalis of Meynert (NBM) in postmortem human brain tissues to investigate neuropathological changes. 71 postmortem NBM samples were grouped by T1DM, T2DM and non-diabetic controls, with Braak stage 0-2 or 3-6. T1DM subjects had only Braak stage 0-2 and were thus compared only to controls with a similar Braak stage and not subjects with Braak stage 3-6. We analysed neurons expressing choline acetyltransferase (ChAT), phosphorylated-Tau, glial cells and vasculature with respective markers. We found significantly less neuronal expression of ChAT in T1DM compared to controls and T2DM with Braak stage 0-2. Later-stage hyperphosphorylated-Tau levels were higher in T2DM compared to controls with Braak stages 3-6. Our results suggest that reduced acetylcholine production by NBM neurons might underlie the cognitive complains of people with T1DM. In contrast, T2DM may exacerbate neuropathological changes associated with Alzheimer’s disease-like alterations.

## Introduction

People with type 1 diabetes (T1DM) or type 2 diabetes (T2DM) often experience cognitive impairment and are both at an increased risk of developing Alzheimer’s disease (AD) [54, 67]. The aetiology of T1DM and T2DM is different: T1DM is caused by an abnormal immune response attacking and destroying the insulin-producing beta cells in the pancreas, and it is often diagnosed at a young age [48]. In contrast, the main driving factors for T2DM are overweight/obesity, sedentary lifestyle, and the consumption of unhealthy diets, characterized by insulin resistance, and typically T2DM develops later in life [68]. These differences are also associated with differences in cognitive impairment, as a previous study found that people with T1DM had more severe cognitive impairment compared with those with T2DM [31]. Yet, despite extensive cognitive assessments, neuroimaging and epidemiological studies [31, 39, 54, 62, 68], the brain mechanisms at the cellular level underlying these cognitive impairments remain unclear, particularly those related to cognitive complains in people with T1DM.

Acetylcholine (ACh)-producing neurons in the cholinergic nucleus basalis of Meynert (NBM) are crucial for cognition. Dysfunction of these neurons leads to cognitive dysfunctions such as memory loss, learning difficulties, mood disorders, and attention deficit [9, 10, 17, 30, 33, 38, 60]. Previous studies have shown that cognitive decline is associated with neuronal atrophy and an increase in hyperphosphorylated tau (p-Tau) in the NBM [45, 69]. Surrounding these neurons, an essential supporting unit is formed by microglia, astrocytes, blood vessels and the glymphatic system. These cells maintain brain homeostasis, ensure neuronal survival, and clear brain waste [15, 20, 22, 36, 55, 59, 64]. In particular, microglia-driven neuroinflammation is known to be pivotal in the pathogenesis of AD [32], whereas studies on the glymphatic system have produced conflicting findings in AD brains [46, 58].

Given these observations, we systemically profiled the neurons in the NBM using markers for choline acetyltransferase (ChAT, the key enzyme that catalyses the biosynthesis of ACh) and Golgi matrix protein GA130 for neuronal activity [6, 9, 24, 57]. We also examined three p-Tau markers (CP13, PHF, AT8) associated with early or later stages of dementia [42], microglia (using ionized calcium binding adaptor molecule 1, Iba1), astrocytes (using glial fibrillary acidic protein, GFAP), vasculature (using alpha-smooth muscle actin, alpha-SMA, a major structural protein expressed by arteries and arterioles [53]), and the glymphatic system (using aquaporin 4, AQP4 [22, 36]). Our goal was to investigate whether the NBM was affected according to these parameters in T1DM or T2DM and to reveal mechanisms linking diabetes to AD.

## Materials and Methods

### Subject information

All brain material was obtained from the Netherlands Brain Bank. The donor or their next of kin gave informed consent for a brain autopsy and for the use of the brain material and medical records for research purposes [29]. In total, 71 postmortem human NBM samples from people with T1DM or T2DM and controls without diabetes were studied (Table 1 & 2). To ensure a fair comparison without the influence of neuropathological changes associated with Alzheimer’s or Parkinson’s disease, we grouped the subjects based on their Braak staging, which reflects the distribution of p-Tau throughout the brain rather than the intensity of p-Tau accumulation [5]. Among all the 10 people with T1DM, Braak stages analysis using AT8 immunohistochemistry (AT8-ir) [5] showed that 9 of them had Braak stages 0-2. Therefore, we sub-grouped all non-diabetic, T1DM and T2DM subjects into two categories: with Braak stage 0-2 (associated with no cognitive decline) or Braak stage 3-6 (associated with mild or severe cognitive decline), [5]. The non-diabatic control groups consisted of 23 subjects, of which 15 subjects with Braak stage 0-2, and 8 with Braak stage 3-6. The T1DM group consisted of 9 subjects with Braak stage 0-2. The T2DM group was composed of 17 subjects with Braak stage 0-2, and 22 subjects with Braak stage 3-6 (Table 1 & 2). In addition, all groups were matched for sex, age, fixation time, post-mortem delay (PMD), pH in cerebrospinal fluid (CSF) (a measure for agonal state) to prevent potential impact of confounding factors. An overview with more detailed information on each individual subject, including APOE sub-genotype, medication use, clinical diagnosis and cause of death, is provided in Table 2.

**Table 1.**
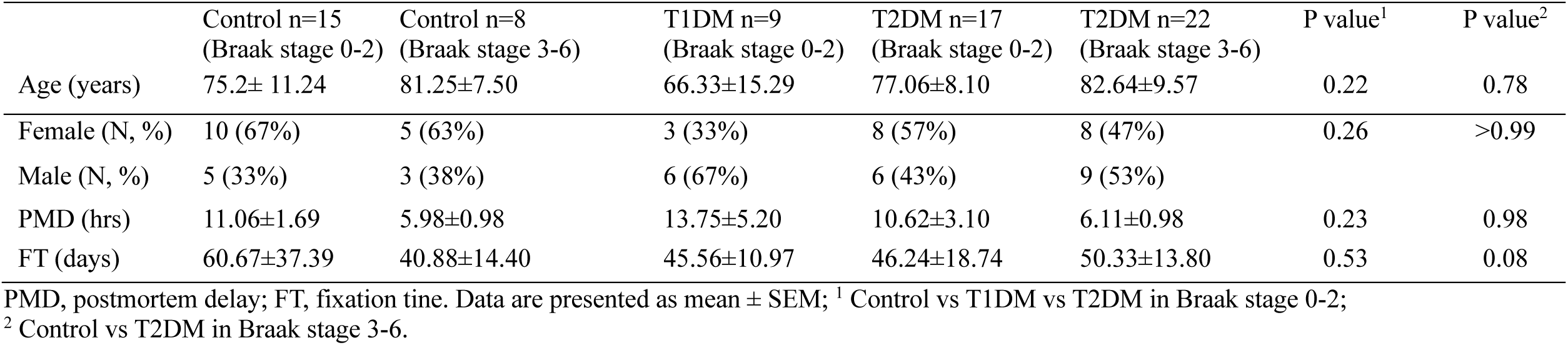
Subjects group characteristics.

**Table 2.**
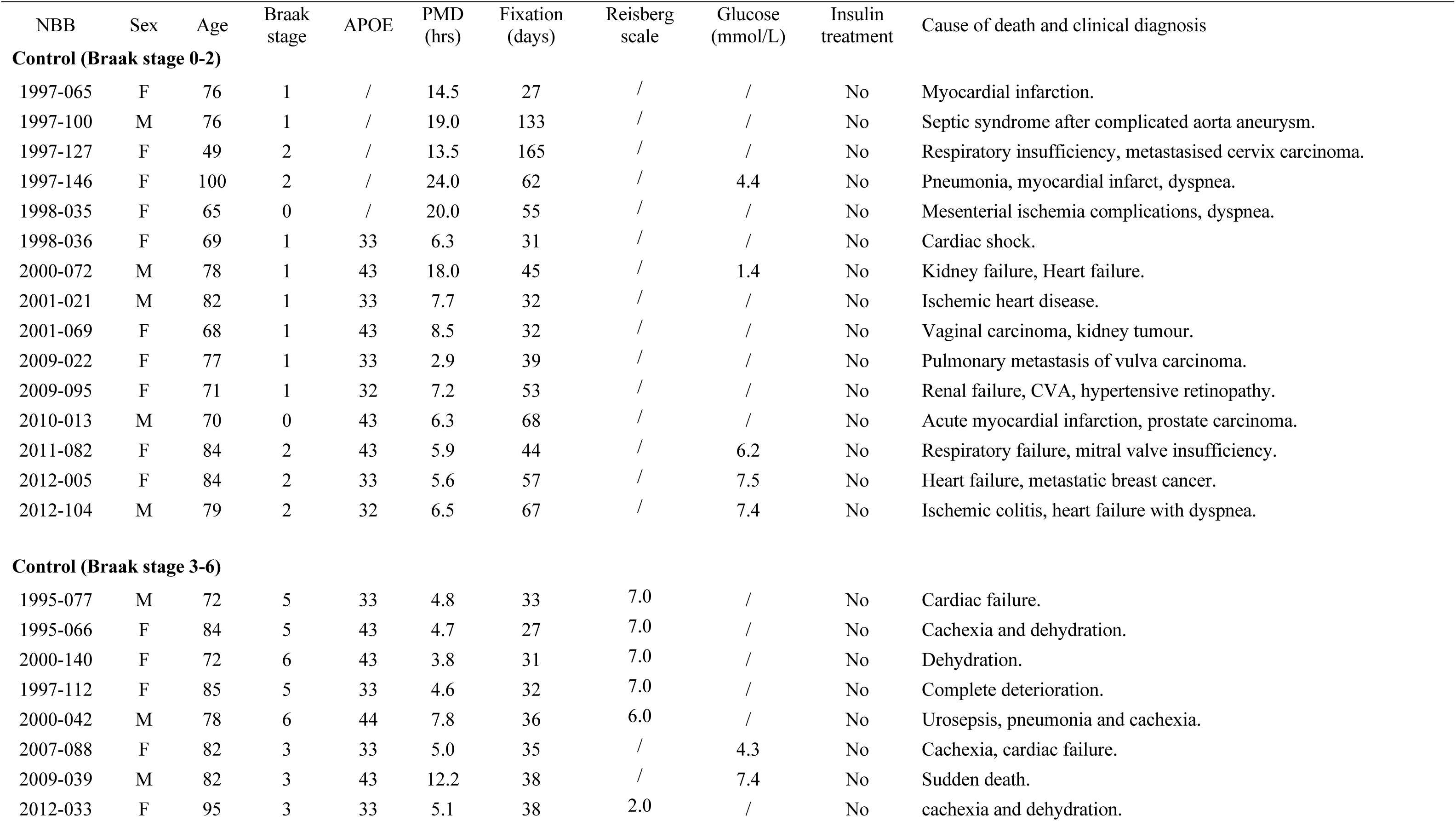
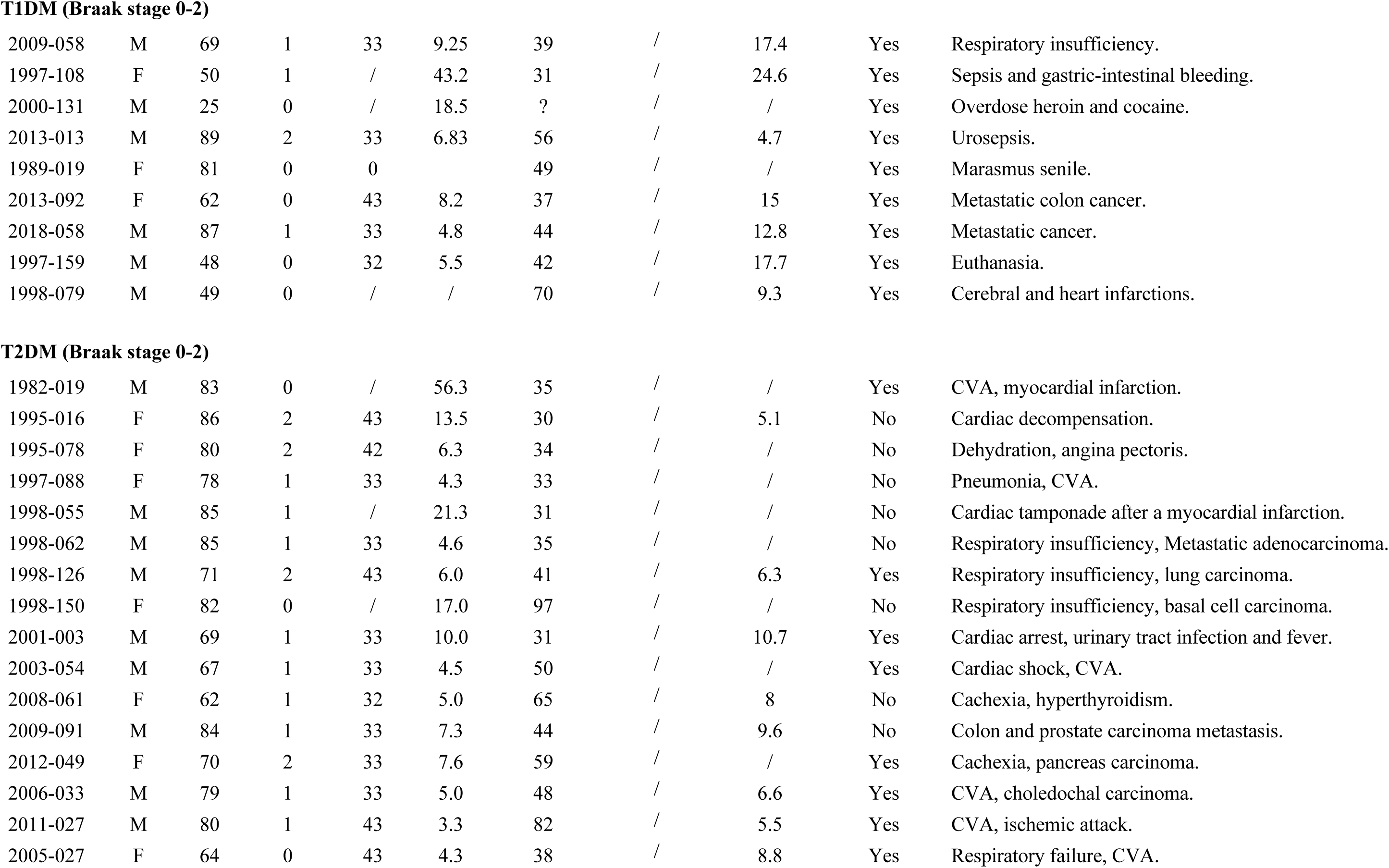
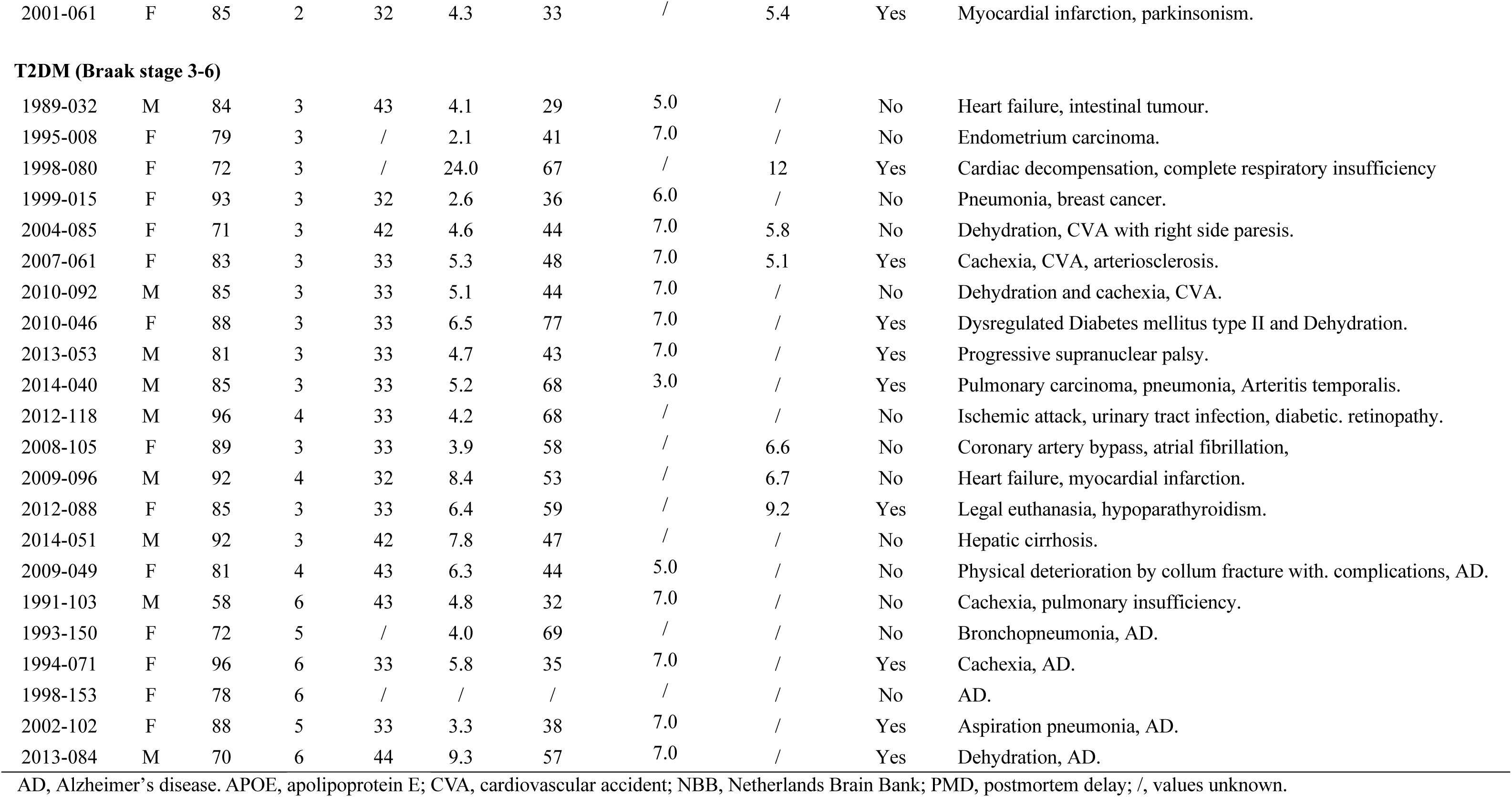
Clinico-pathological details of subjects.

| NBB | Sex | Age | Braak stage | APOE | PMD (hrs) | Fixation (days) | Reisberg scale | Glucose (mmol/L) | Insulin treatment | Cause of death and clinical diagnosis |
| --- | --- | --- | --- | --- | --- | --- | --- | --- | --- | --- |
| **Control (Braak stage 0-2)** |  |  |  |  |  |  |  |  |  |  |
| 1997-065 | F | 76 | 1 | / | 14.5 | 27 | / | / | No | Myocardial infarction. |
| 1997-100 | M | 76 | 1 | / | 19.0 | 133 | / | / | No | Septic syndrome after complicated aorta aneurysm. |
| 1997-127 | F | 49 | 2 | / | 13.5 | 165 | / | / | No | Respiratory insufficiency, metastasised cervix carcinoma. |
| 1997-146 | F | 100 | 2 | / | 24.0 | 62 | / | 4.4 | No | Pneumonia, myocardial infarct, dyspnea. |
| 1998-035 | F | 65 | 0 | / | 20.0 | 55 | / | / | No | Mesenterial ischemia complications, dyspnea. |
| 1998-036 | F | 69 | 1 | 33 | 6.3 | 31 | / | / | No | Cardiac shock. |
| 2000-072 | M | 78 | 1 | 43 | 18.0 | 45 | / | 1.4 | No | Kidney failure, Heart failure. |
| 2001-021 | M | 82 | 1 | 33 | 7.7 | 32 | / | / | No | Ischemic heart disease. |
| 2001-069 | F | 68 | 1 | 43 | 8.5 | 32 | / | / | No | Vaginal carcinoma, kidney tumour. |
| 2009-022 | F | 77 | 1 | 33 | 2.9 | 39 | / | / | No | Pulmonary metastasis of vulva carcinoma. |
| 2009-095 | F | 71 | 1 | 32 | 7.2 | 53 | / | / | No | Renal failure, CVA, hypertensive retinopathy. |
| 2010-013 | M | 70 | 0 | 43 | 6.3 | 68 | / | / | No | Acute myocardial infarction, prostate carcinoma. |
| 2011-082 | F | 84 | 2 | 43 | 5.9 | 44 | / | 6.2 | No | Respiratory failure, mitral valve insufficiency. |
| 2012-005 | F | 84 | 2 | 33 | 5.6 | 57 | / | 7.5 | No | Heart failure, metastatic breast cancer. |
| 2012-104 | M | 79 | 2 | 32 | 6.5 | 67 | / | 7.4 | No | Ischemic colitis, heart failure with dyspnea. |
| **Control (Braak stage 3-6)** |  |  |  |  |  |  |  |  |  |  |
| 1995-077 | M | 72 | 5 | 33 | 4.8 | 33 | 7.0 | / | No | Cardiac failure. |
| 1995-066 | F | 84 | 5 | 43 | 4.7 | 27 | 7.0 | / | No | Cachexia and dehydration. |
| 2000-140 | F | 72 | 6 | 43 | 3.8 | 31 | 7.0 | / | No | Dehydration. |
| 1997-112 | F | 85 | 5 | 33 | 4.6 | 32 | 7.0 | / | No | Complete deterioration. |
| 2000-042 | M | 78 | 6 | 44 | 7.8 | 36 | 6.0 | / | No | Urosepsis, pneumonia and cachexia. |
| 2007-088 | F | 82 | 3 | 33 | 5.0 | 35 | / | 4.3 | No | Cachexia, cardiac failure. |
| 2009-039 | M | 82 | 3 | 43 | 12.2 | 38 | / | 7.4 | No | Sudden death. |
| 2012-033 | F | 95 | 3 | 33 | 5.1 | 38 | 2.0 | / | No | cachexia and dehydration. |
| **T1DM (Braak stage 0-2)** |  |  |  |  |  |  |  |  |  |  |
| 2009-058 | M | 69 | 1 | 33 | 9.25 | 39 | / | 17.4 | Yes | Respiratory insufficiency. |
| 1997-108 | F | 50 | 1 | / | 43.2 | 31 | / | 24.6 | Yes | Sepsis and gastric-intestinal bleeding. |
| 2000-131 | M | 25 | 0 | / | 18.5 | ? | / | / | Yes | Overdose heroin and cocaine. |
| 2013-013 | M | 89 | 2 | 33 | 6.83 | 56 | / | 4.7 | Yes | Urosepsis. |
| 1989-019 | F | 81 | 0 | 0 |  | 49 | / | / | Yes | Marasmus senile. |
| 2013-092 | F | 62 | 0 | 43 | 8.2 | 37 | / | 15 | Yes | Metastatic colon cancer. |
| 2018-058 | M | 87 | 1 | 33 | 4.8 | 44 | / | 12.8 | Yes | Metastatic cancer. |
| 1997-159 | M | 48 | 0 | 32 | 5.5 | 42 | / | 17.7 | Yes | Euthanasia. |
| 1998-079 | M | 49 | 0 | / | / | 70 | / | 9.3 | Yes | Cerebral and heart infarctions. |
| **T2DM (Braak stage 0-2)** |  |  |  |  |  |  |  |  |  |  |
| 1982-019 | M | 83 | 0 | / | 56.3 | 35 | / | / | Yes | CVA, myocardial infarction. |
| 1995-016 | F | 86 | 2 | 43 | 13.5 | 30 | / | 5.1 | No | Cardiac decompensation. |
| 1995-078 | F | 80 | 2 | 42 | 6.3 | 34 | / | / | No | Dehydration, angina pectoris. |
| 1997-088 | F | 78 | 1 | 33 | 4.3 | 33 | / | / | No | Pneumonia, CVA. |
| 1998-055 | M | 85 | 1 | / | 21.3 | 31 | / | / | No | Cardiac tamponade after a myocardial infarction. |
| 1998-062 | M | 85 | 1 | 33 | 4.6 | 35 | / | / | No | Respiratory insufficiency, Metastatic adenocarcinoma. |
| 1998-126 | M | 71 | 2 | 43 | 6.0 | 41 | / | 6.3 | Yes | Respiratory insufficiency, lung carcinoma. |
| 1998-150 | F | 82 | 0 | / | 17.0 | 97 | / | / | No | Respiratory insufficiency, basal cell carcinoma. |
| 2001-003 | M | 69 | 1 | 33 | 10.0 | 31 | / | 10.7 | Yes | Cardiac arrest, urinary tract infection and fever. |
| 2003-054 | M | 67 | 1 | 33 | 4.5 | 50 | / | / | Yes | Cardiac shock, CVA. |
| 2008-061 | F | 62 | 1 | 32 | 5.0 | 65 | / | 8 | No | Cachexia, hyperthyroidism. |
| 2009-091 | M | 84 | 1 | 33 | 7.3 | 44 | / | 9.6 | No | Colon and prostate carcinoma metastasis. |
| 2012-049 | F | 70 | 2 | 33 | 7.6 | 59 | / | / | Yes | Cachexia, pancreas carcinoma. |
| 2006-033 | M | 79 | 1 | 33 | 5.0 | 48 | / | 6.6 | Yes | CVA, choledochal carcinoma. |
| 2011-027 | M | 80 | 1 | 43 | 3.3 | 82 | / | 5.5 | Yes | CVA, ischemic attack. |
| 2005-027 | F | 64 | 0 | 43 | 4.3 | 38 | / | 8.8 | Yes | Respiratory failure, CVA. |
| 2001-061 | F | 85 | 2 | 32 | 4.3 | 33 | / | 5.4 | Yes | Myocardial infarction, parkinsonism. |
| **T2DM (Braak stage 3-6)** |  |  |  |  |  |  |  |  |  |  |
| 1989-032 | M | 84 | 3 | 43 | 4.1 | 29 | 5.0 | / | No | Heart failure, intestinal tumour. |
| 1995-008 | F | 79 | 3 | / | 2.1 | 41 | 7.0 | / | No | Endometrium carcinoma. |
| 1998-080 | F | 72 | 3 | / | 24.0 | 67 | / | 12 | Yes | Cardiac decompensation, complete respiratory insufficiency |
| 1999-015 | F | 93 | 3 | 32 | 2.6 | 36 | 6.0 | / | No | Pneumonia, breast cancer. |
| 2004-085 | F | 71 | 3 | 42 | 4.6 | 44 | 7.0 | 5.8 | No | Dehydration, CVA with right side paresis. |
| 2007-061 | F | 83 | 3 | 33 | 5.3 | 48 | 7.0 | 5.1 | Yes | Cachexia, CVA, arteriosclerosis. |
| 2010-092 | M | 85 | 3 | 33 | 5.1 | 44 | 7.0 | / | No | Dehydration and cachexia, CVA. |
| 2010-046 | F | 88 | 3 | 33 | 6.5 | 77 | 7.0 | / | Yes | Dysregulated Diabetes mellitus type II and Dehydration. |
| 2013-053 | M | 81 | 3 | 33 | 4.7 | 43 | 7.0 | / | Yes | Progressive supranuclear palsy. |
| 2014-040 | M | 85 | 3 | 33 | 5.2 | 68 | 3.0 | / | Yes | Pulmonary carcinoma, pneumonia, Arteritis temporalis. |
| 2012-118 | M | 96 | 4 | 33 | 4.2 | 68 | / | / | No | Ischemic attack, urinary tract infection, diabetic. retinopathy. |
| 2008-105 | F | 89 | 3 | 33 | 3.9 | 58 | / | 6.6 | No | Coronary artery bypass, atrial fibrillation, |
| 2009-096 | M | 92 | 4 | 32 | 8.4 | 53 | / | 6.7 | No | Heart failure, myocardial infarction. |
| 2012-088 | F | 85 | 3 | 33 | 6.4 | 59 | / | 9.2 | Yes | Legal euthanasia, hypoparathyroidism. |
| 2014-051 | M | 92 | 3 | 42 | 7.8 | 47 | / | / | No | Hepatic cirrhosis. |
| 2009-049 | F | 81 | 4 | 43 | 6.3 | 44 | 5.0 | / | No | Physical deterioration by collum fracture with. complications, AD. |
| 1991-103 | M | 58 | 6 | 43 | 4.8 | 32 | 7.0 | / | No | Cachexia, pulmonary insufficiency. |
| 1993-150 | F | 72 | 5 | / | 4.0 | 69 | / | / | No | Bronchopneumonia, AD. |
| 1994-071 | F | 96 | 6 | 33 | 5.8 | 35 | 7.0 | / | Yes | Cachexia, AD. |
| 1998-153 | F | 78 | 6 | / | / | / | / | / | No | AD. |
| 2002-102 | F | 88 | 5 | 33 | 3.3 | 38 | 7.0 | / | Yes | Aspiration pneumonia, AD. |
| 2013-084 | M | 70 | 6 | 44 | 9.3 | 57 | 7.0 | / | Yes | Dehydration, AD. |

AD, Alzheimer's disease. APOE, apolipoprotein E; CVA, cardiovascular accident; NBB, Netherlands Brain Bank; PMD, postmortem delay; /, values unknown.

### Histology and morphometry of the Nucleus Basalis of Meynert

After autopsy, the isolated brain tissues containing the hypothalamus and NBM were immediately immersed in formalin and fixed at room temperature for 1-2 months. The tissues were then ethanol-dehydrated, toluene-cleared and paraffin-embedded. Serial 6 μm serial coronal sections were made along the rostro-caudal axis including the NBM area. The anatomical orientation and rostro-caudal range of the NBM were determined by Nissl staining (Supplementary information) and confirmed by ChAT immunoreactivity (ChAT-ir) (see 2.3). Since the NBM consists of a large cell population, we performed our studies in a standardized region of the NBM, as described in previous studies [9, 37, 69], specifically the anteromedial part (Ch4am) and the anterolateral part (Ch4al) of the NBM at the level of the fornix and/or the anterior commissure [60] (Fig.1a). All immunohistochemical or immunofluorescent staining were performed on consecutive sections at this level. For ChAT-ir, three sections per brain were selected: the level with peak neuron density in the NBM and sections 600 μm (100 sections) anterior and posterior to it. For other staining, one section per subject at the level of peak neuron density in the NBM was used.

### Immunohistochemistry and Immunofluorescence

For antibodies of ChAT, GA130, CP13, PHF1, Iba1, GFAP and alpha-SMA, we performed epitope retrieval to enhance antigen exposure (Supplementary information). AT8, AQP4 antibody did not require epitope retrieval. After these procedures, sections were incubated with the primary antibody diluted in SUMI buffer (0.25% gelatine, 0.5% Triton X-100 in TBS (pH 7.6), except for the alpha-SMA and GFAP antibodies, which were diluted in SUMI containing 10% donkey serum. All primary antibodies were incubated overnight at 4℃. For immunohistochemistry, after overnight incubation, sections were incubated with corresponding biotinylated secondary antibody followed by avidin-biotin complex (ABC). The chromogenic reaction was then performed with 3,3’-Diaminobenzidine with or without ammonium nickel sulphate (Supplementary information). All reactions were stopped in distilled water, ethanol-dehydrated, xylene-cleared and cover-slipped with Entellan mounting medium. For immunofluorescence, sections were incubated with the corresponding biotinylated secondary antibody followed by a streptavidin fluorescence antibody, and nuclei were labelled with DAPI. The list of primary antibodies and information on their specificity is shown in Supplementary Table 1.

### Imaging acquisition and quantitative analysis

All images, except for AQP4-ir, were acquired using an Axio Scanner (Zeiss) and analyzed with QuPath. An outline of the NBM was manually drawn at 20x magnification, and the threshold for a positive signal of the mask was set at two times the optical density (O.D.) of the background (O.D. was used to express all data, as the main conclusions remain the same when integrated optical density (IOD = O.D. x % area) is used as an endpoint.) Total neuron density was calculated based on total Nissl-stained cell count and area of the NBM. The number of immunoreactive areas (ir area), representing the locations of individual neurons, glial cells and blood vessels, was quantified in the NBM region, which was delineated based on Nissl staining. The average number of immunoreactive areas was calculated using the ‘Intensity Features’ tool in QuPath software and normalized to the background. For AQP4-ir, images were captured using Image Pro version 6.3 (Media Cybernetics), and the relative area of the positive signal was quantified using the “Analyze Particles” tool in ImageJ. The term “particles” refers to Iba1-ir microglial soma, with coverage areas ranging from 20 µm2 to 100 µm2, as defined in a previous study [26]. The relative area of coverage for alpha-SMA-ir and GFAP-ir was measured using the “pixel classification” tool in QuPath.

### Statistical analysis

Following the normality test (D’Agostino and Pearson test), pairwise comparisons within the Braak stage 0–2 group or the Braak stage 3–6 group were performed using Mann-Whitney U tests. Comparisons between control subjects with Braak stage 0–2 or Braak stage 3–6 and T2DM subjects with Braak stage 3–6 were also conducted using Mann-Whitney U tests. To identify differences among the three groups, the Kruskal-Wallis test was applied due to the non-normal distribution of the data. Correlations and confounder analyses were evaluated using Spearman’s rank correlation coefficient. A p-value ≤ 0.05 was considered statistically significant. All statistical tests were performed using GraphPad Prism 9.5.1.

### Confounder analyses

Considering the potential impact of biological and pathological variabilities in the human subjects before and during death, including age, postmortem delay (PMD) affecting tissue degradation, pH of CSF, blood glucose levels during lifetime, tissue fixation time, and Braak stage, we ensured maximum matching of these parameters across all groups. Furthermore, we have conducted confounder analyses between these factors and the major study outcomes.

## Results

### Diverse changes of cholinergic neurons in the NBM of individuals with T1DM or T2DM

We first quantified the density of NBM neurons per delineated NBM area using Nissl staining, soma size between 30 µm^2^ - 300 µm^2^ was defined as a neuron. Neuron density of control, T1DM, and T2DM subjects across different Braak stages was comparable (Fig. 1b, c). Moreover, in confounder analysis, we found no corelations between neuron density and Age, PMD, pH of CSF, tissue fixation time, blood glucose level or Braak stage (Fig. S1). However, in the NBM of T1DM subjects, we found significantly a lower ChAT immunoreactive (ChAT-ir) area and optical density (O.D.) of ChAT-ir. This was not observed in controls and T2DM subjects with Braak stage 0-2 (Figs. 1d-f). In controls with Braak stage 3-6, we observed a significantly lower optical density of ChAT compared to controls with Braak stage 0-2. Interestingly, in T2DM subjects with Braak stage 3-6, the ChAT-ir area was not smaller than in controls and T2DM subjects with Braak stage 0-2. Moreover, the ChAT-ir optical density was significantly higher in T2DM subjects with Braak stages 3-6 than in controls of the same Braak stage group (Figs. 1d-f). These results indicate that cognitive impairments in patients with T1DM are likely associated with cholinergic neuronal dysfunction in the NBM; however, this seems be not the case in patients with T2DM.

**Fig. 1.**
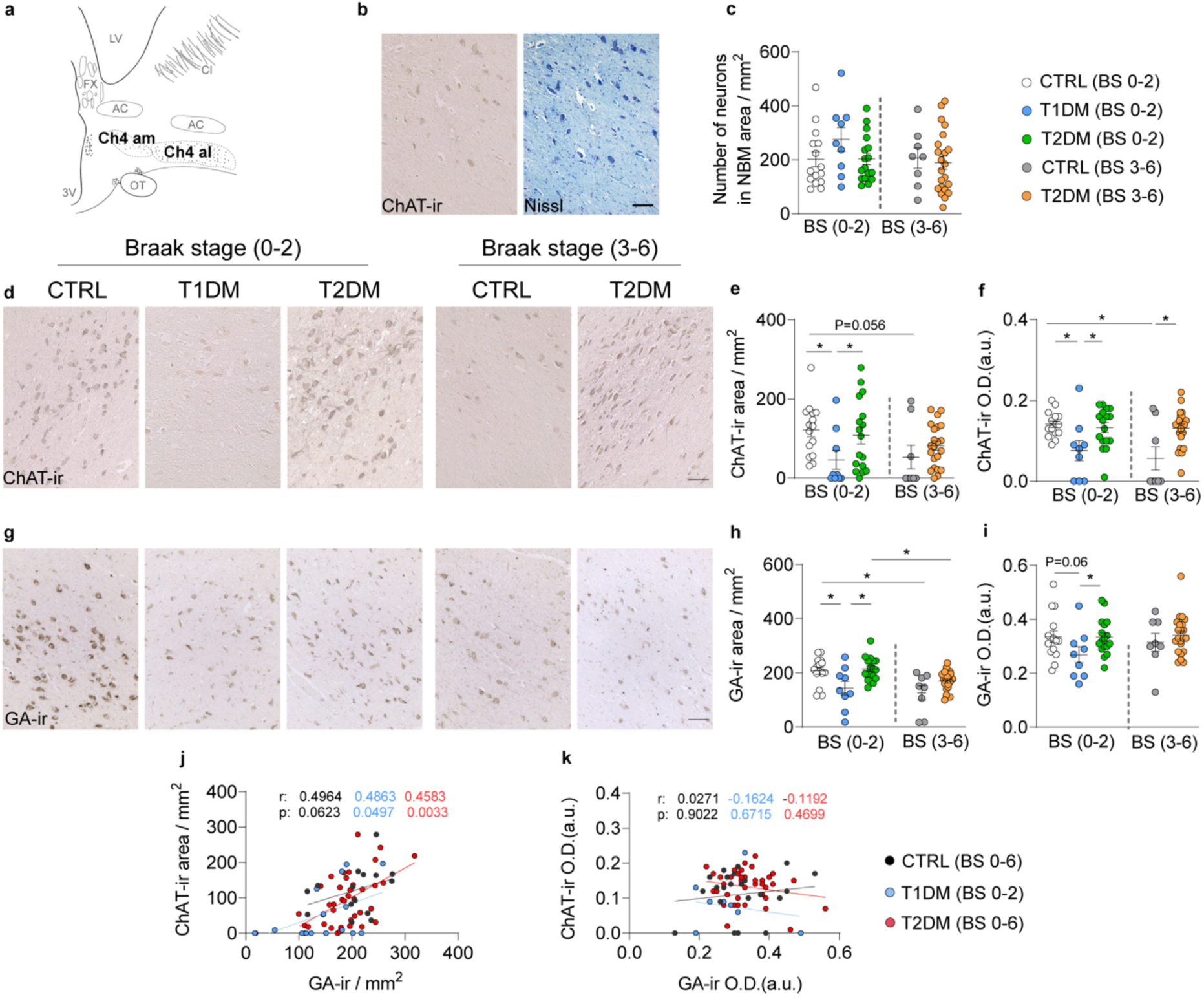
Reductions of choline acetyltransferase immunoreactive (ChAT-ir) and Golgi matrix protein GA130 immunoreactive (GA-ir) neurons in the NBM of T1DM subjects. **a** Schematic representation of the medial part (Ch4-am) and the lateral part (Ch4-al) of the nucleus basalis of Meynert (NBM) area used for cell profiling. 3V, third ventricle; AC, anterior commissure; CI, capsula interna; FX, fornix; LV, lateral ventricle; ON, optical nerve. **b, c** Neuron density visualized by Nissl staining in the NBM of control, T1DM, and T2DM subjects over different Braak stages is comparable. **d** Representative images of ChAT-ir neurons and Nissl-stained cells in T1DM subjects. Representative images of ChAT-ir in the NBM of the control, T1DM and T2DM subjects. **e, f** Comparisons of ChAT-ir area and optical density (in arbitrary units, O.D. (a.u.)) of ChAT-ir neurons between controls, T1DM and T2DM with Braak stage (BS) 0-2 or 3-6. g Representative images of GA-ir in the NBM of the controls, T1DM and T2DM subjects. **h, i** Comparisons of GA-ir area and O.D. of GA-ir neurons between controls, T1DM and T2DM with BS 0-2 or BS 3-6. **j, k** Plots of GA-ir area and O.D. with ChAT-ir O.D. Scale bar: 100 µm in B, D & G. Data are represented as mean ± SEM. * p<0.05.

As no clear difference was found in cell density in the NBM by Nissl histology, we wondered whether the fewer ChAT-ir in the NBM was due to less neuronal activity Therefore, we analysed the metabolic activity of the NBM using GA130, which recognizes the Golgi matrix and has been validated as an indicator of neuronal activity in the NBM [9, 49]. We found that the GA-ir area and optical density were indeed lower in the NBM of T1DM subjects compared to controls and T2DM subjects (Figs. 1g-i). Moreover, consistent with previous findings [9, 49], the optical density of GA-ir in controls and T2DM subjects with Braak stages 3-6 were both significantly lower than those with Braak stages 0-2 (Figs. 1g-i). Importantly, we found a positive correlation between the GA-ir area and the ChAT-ir area in T1DM and T2DM subjects, while a trend was present in the controls (Figs. 1j, k), suggesting that reduced cellular metabolic activity might underlie the decreased ACh production in these neurons in T1DM individuals.

With respect to the major confounders (Fig. S2; S3), we found a negative correlation between ChAT-ir areas with Braak stage in control subjects (Fig. S2f), and a negative correlation between the GA-ir areas and Braak stage in control and T2DM subjects (Fig. S3f), indicating the NBM neuronal activity declines along the course of cognitive decline.

### The impact of T2DM on P-Tau in the NBM

Since all the T1DM subjects included in our analysis had Braak stage 0-2, as determined by AT8, an indicator of later stage of AD, we investigated whether T1DM brains expressed early-stage p-Tau. We examined two additional p-Tau proteins, the CP13, which appears at a relatively early stage (pre-tangle stage) of AD [43], and the PHF1, which appears at a later stage of cognitive decline [43]. In subjects with Braak stage 0-2, we found no difference in the expression of all three p-Tau proteins between control, T1DM and T2DM subjects (Fig. 2). Interestingly, in subjects with Braak stage 3-6, the early-stage p-Tau CP13-ir areas was significantly lower in T2DM subjects compared to controls (Figs. 2a-i). These findings suggest that T2DM may exacerbate neuropathological changes associated with AD compared to non-diabetic individuals.

**Fig. 2.**
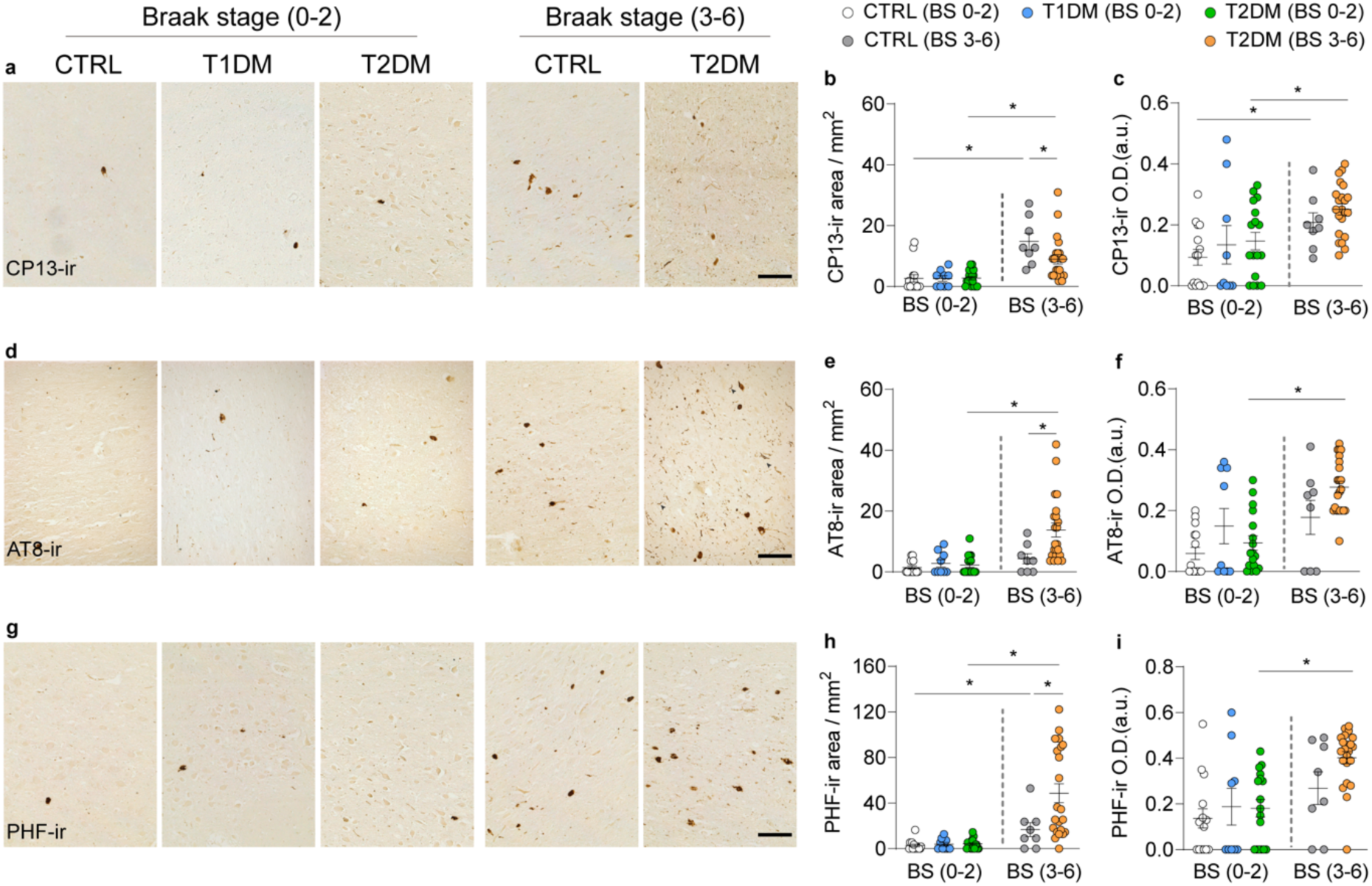
P-Tau did not change in T1DM but increased in T2DM subjects with Braak stage 3-6. **a** Representative image of CP13-immunoreactive (Cp13-ir) neurons in the NBM of control, T1DM and T2DM subjects. **b, c** Comparisons of the CP13-ir area and CP13-ir optical density (in arbitrary units, O.D. (a.u)) between controls, T1DM and T2DM with Braak stage (BS) 0-2 or 3-6. **d** Representative images of AT8-ir neurons in the NBM of control, T1DM and T2DM subjects. Note that the AT8-ir was also present in neurites. **e, f** Comparisons of AT8-ir area and AT8-ir O.D. neurons between control, T1DM and T2DM with BS 0-2 or BS 3-6. **g** Representative images of PHF-ir neurons in the NBM of the control, T1DM and T2DM subjects. **h, i** Comparisons of PHF-ir area and PHF-ir O.D. between control, T1DM and T2DM in BS 0-2 or BS 3-6. Scale bar: 100 µm in (a, d, g). Data are represented as mean ± SEM. * p<0.05

Regarding the impact of major confounders on p-Tau expression (Figs. S4; S5; S6), all three p-Tau markers showed a positive correlation between the immunoreactive areas or the optic density and the Braak stage in both control and T2DM subjects, indicating that p-Tau levels increase as cognitive decline progresses (Figs. S4f, l; S5f, l; S6f, l). Additionally, CP13-ir and PHF-ir areas were positively associated with age in controls (Fig. S4a; Fig. S6a). Furthermore, a negative correlation was observed between ChAT-ir and CP13-ir areas in the control group (Figs. S4m, 4n). None of the confounders showed a correlation with p-Tau expression in the NBM of T1DM subjects.

### Lower microglial activity in the NBM of T1DM individuals

Microglia-driven neuroinflammation is recognized as pivotal in the pathogenesis of AD [32]. However, among all the microglial parameters, *i.e.* the soma density, masked area (%) and the soma size of Iba1-ir, we only observed a decrease in Iba1-ir microglial soma density in the NBM of T1DM subjects with Braak stages 0-2 (Figs. 3a, 3b). No significant changes were observed in microglial cell density, soma size, or total Iba1-ir area in controls and T2DM subjects with Braak stages 3-6. Additionally, during confounder analysis (Figs. S7; S8), we found a negative correlation between Iba1-ir soma size and Braak stage in the control group (Fig. S7r). Interestingly, we found a negative correlation between the Iba1-ir soma size with the of ChAT-ir areas and optical density in the controls without diabetes (Figs. S8c, f) and a positive correlation between Iba1-ir cell density and ChAT-ir optical density, in both controls and T2DM (Fig. S8d) suggesting microglial activity might be associated with cholinergic neuronal activity in the NBM.

**Fig. 3.**
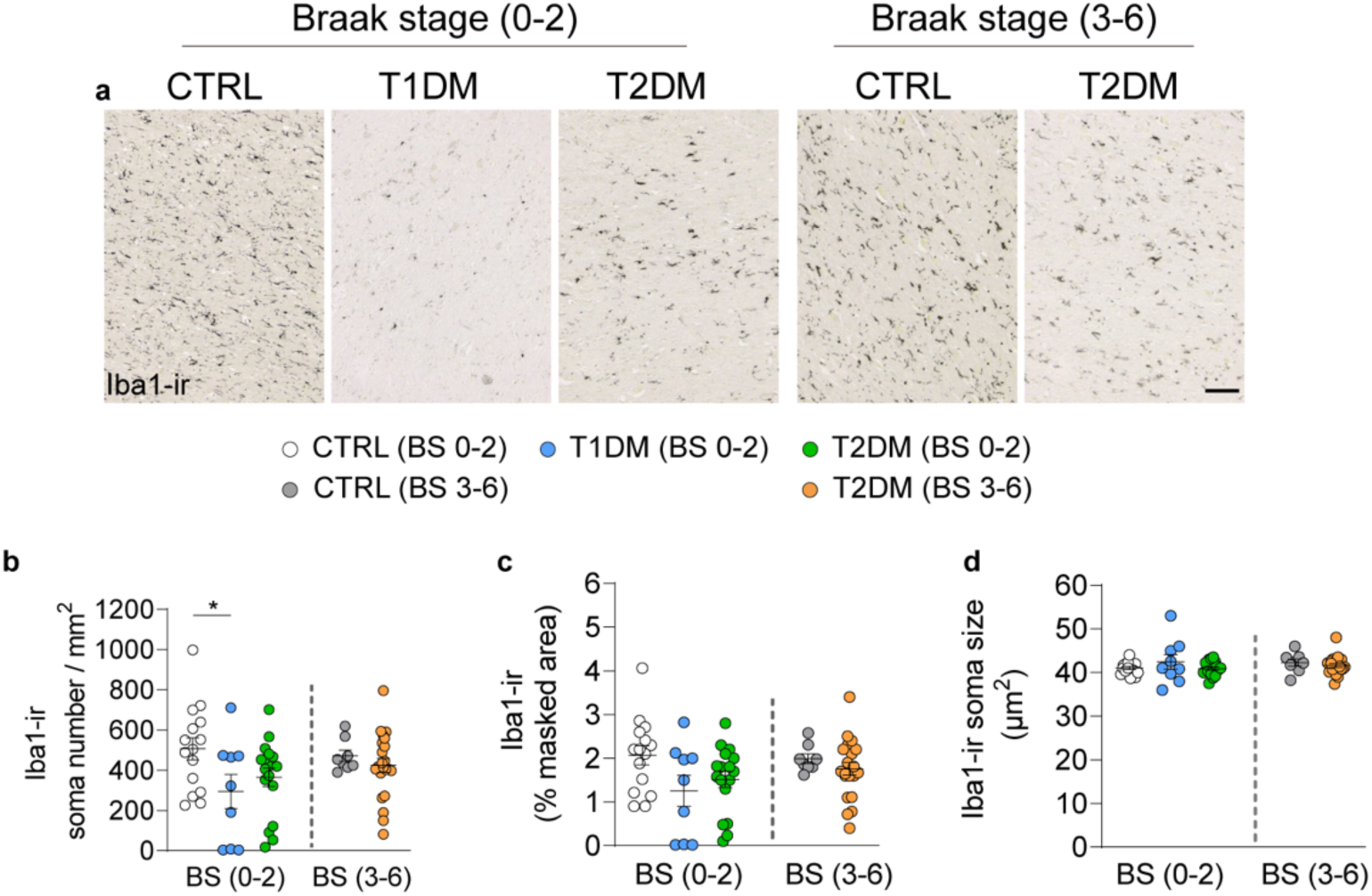
Reduced Iba1-ir microglia in the NBM of T1DM subjects. **a** Representative images of ionized calcium binding adaptor molecule 1 immunoreactive (Iba1-ir) in the NBM of control, T1DM and T2DM subjects. **b, c, d** Comparisons of the soma density, masked area (%) and the soma size of Iba1-ir between control, T1DM and T2DM with Braak stage (BS) 0-2 or 3-6. Note that Iba1-ir is lower in both T1DM and T2DM subjects with BS 0-2. Scale bar: 100 µm. Data are represented as mean ± SEM.* p<0.05.

### The impact of diabetes on astroglia and the glymphatic system

In addition to microglia, astrocytes also play a significant role in AD pathogenesis [1, 14]. We found both T1DM and T2DM with Braak stage 0-2 had lower GFAP-ir astrocytes compared to controls with Braak stage 0-2. Moreover, controls with Braak stage 3-6 showed reduced numbers of GFAP-ir astrocytes compared to the controls with Braak stage 0-2. In contrast, no difference was observed between T2DM subjects with Braak stages 0-2 and those with Braak stages 3-6 (Figs. 4a, b). These findings suggest that GFAP-ir astrocytes are affected by diabetes- or AD-associated neuropathology. A subpopulation of astrocytes that express AQP4 is a key component of the brain glymphatic system which is crucial for clearing brain waste, including amyloid-β [22, 36]. Previous studies have reported conflicting results on AQP4 levels in AD brains [46, 58]. Interestingly, we found that AQP4-ir was lower in T1DM compared to controls and T2DM subjects (Figs. 4c - e). Conversely, AQP4-ir in controls with low or high Braak stages did not show differences, suggesting that AQP4 levels in the NBM are not associated with AD pathology. However, during confounder analysis (Fig. S9), we did find a positive correlation between AQP4-ir and ChAT-ir in controls (Figs. S9g, h), indicating that astrocytic dysfunction might be associated with a loss of ChAT-ir in the NBM.

**Fig. 4.**
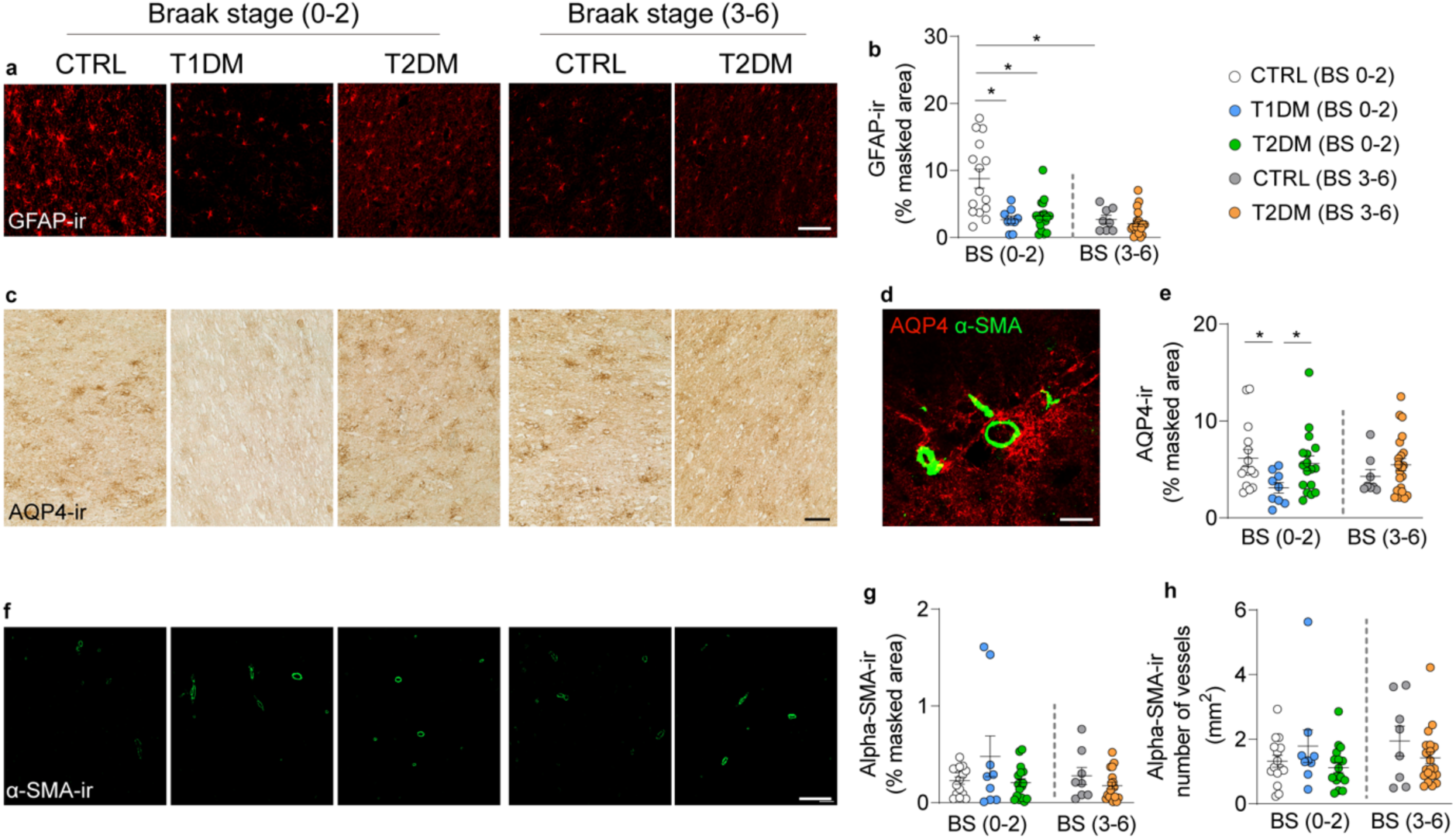
Different impact of diabetes on astroglia and glymphatic system. **a** Representative images of glial fibrillary acidic protein immunoreactive (GFAP-ir) in the NBM of control, T1DM and T2DM subjects. **b** Comparisons of the masked area (%) of GFAP-ir between control, T1DM and T2DM with Braak stage (BS) 0-2 or 3-6. Note that GFAP-ir is lower in both T1DM and T2DM subjects with BS 0-2 and in controls with BS 3-6. **c** Representative images of aquaporin 4 immunoreactive (AQP4-ir) in the NBM of control, T1DM and T2DM subjects. **d** Illustration of AQP4-ir astrocytes surrounding alpha-SMA-ir vessels that form the peri-vascular glymphatic system. **e** Comparisons of the masked area of AQP4-ir between control, T1DM and T2DM in BS 0-2 or BS 3-6. **f** Representative images of alpha-smooth muscle actin immunoreactive (α-SMA-ir) in the NBM of the control, T1DM and T2DM subjects. **g, h** Comparisons of the alpha-SMA-ir masked area and the number of stained vessels between control, T1DM and T2DM with BS 0-2 or 3-6. Note that AQP4-ir is specifically lower in T1DM subjects. Scale bar: 100 µm in a, 50 µm in c, 30 µm in d, 20 µm in f. Data are represented as mean ± SEM. * p<0.05.

Given the close association between GFAP- and AQP4-expressing astrocytes and the vasculature, we also evaluated alpha-SMA, an endothelial marker for arteries and arterioles. A previous study has found increased alpha-SMA-ir vessels in the hypothalamus of T2DM individuals [66]. However, we found no difference between controls, T1DM and T2DM with different Braak stages (Figs. 4f - h). Among all confounders (Fig. S10; S11), a positive correlation was identified between GFAP-ir and glucose levels in the T1DM group (Fig. S10d). Additionally, alpha-SMA-ir masked area correlated negatively with glucose levels in the same group (Figure S11j).

### Confounder analysis

In addition to the aforementioned factors in confounder analyses (Figs S1 - S11), we found a positive correlation between ChAT-ir and PMD in the control group and with CSF pH value in the T1DM group (Figs. S2b, 2c). There was a negative association between CP13-ir and PMD in the control group (Figure S4B) and a negative correlation of AT8 with fixation time and PMD in the T2DM group (Fig. S5k). Additionally, we observed correlations between AT8 and PMD in controls (Figs. S7b, 7h, 7n) and CSF pH in T2DM (Figs. S7c, 7i). Moreover, there was a positive correlation between Iba1-ir soma size and Braak stage in control subjects (Fig. S7) and a negative correlation between GFAP-ir masked area and Braak stage in both control and T2DM subjects (Fig. S10), indicating that cognitive decline is associated with neuroinflammatory changes in glial cells in the NBM. Further incidental significance for several parameters is shown in Figures S1-S11. Nevertheless, these correlations did not affect our results because the groups were matched for these factors.

## Discussion

To investigate the association between diabetes and cognitive decline, we studied neurons, glial cells and vasculature in the NBM using postmortem brain tissues donated by people with T1DM or T2DM, and those without diabetes as matched controls. We found significantly less ChAT-ir in the NBM of people with T1DM, which correlated with reduced Golgi apparatus matrix protein GA130-ir in NBM neurons, as well as diminished Iba1-ir microglia and AQP4-expressing astrocytes in the same region. This discovery highlights a unique neuron-glia dysfunction associated with T1DM, indicating diminished activity of the NBM and potentially reduced production of ACh underpinning the severe cognitive impairment observed in these individuals. Furthermore, our data showed that among individuals with Braak stage 3-6, those with T2DM exhibited more p-Tau-ir in the NBM compared to the non-diabetic controls, suggesting that T2DM may exacerbate neuropathological changes associated with AD.

Regarding the T1DM pathology underlying the pronounced cholinergic neuronal dysfunction, one of the most plausible explanations is the dark side of insulin - hypoglycemia. Compared to T2DM, people with T1DM experience hypoglycemia more frequently. It is estimated to have a prevalence of 50% in T1DM compared to 10% in T2DM [40]. Frequent hypoglycemia results in larger glucose variability, characterized by fluctuations between hyperglycemic and hypoglycemic states [52]. The detrimental impact of hypoglycemia on cholinergic neurons includes disruption of ATP production and mitochondrial function [28], as well as impaired biosynthesis of ACh from the glucose metabolite Acetyl-CoA. These changes can collectively compromise cholinergic neuronal function. Conversely, cognitive impairment can worsen poor glycemic control because it heavily relies on self-management of the person with diabetes [25, 35, 50], thus creating a detrimental cycle that further impairs brain function. Our results find a decreased trend of the ChAT-ir area in control with Braak stage 3-6 which is similar as found in a previous study. [13]

Insulin is known for its neuroprotective role [2, 19], and has been explored as a potential therapy for AD [11]. However, our results suggest that lifelong insulin treatment in T1DM does not prevent cholinergic neuronal dysfunction in the NBM. A potential explanation is that synthetic insulin supplementation in T1DM may inadequately reach brain cells due to insulin resistance [44, 65]. The blood-brain barrier can limit insulin transport into the CNS during the circumstances of hyperinsulinemia or insulin resistance [27, 47], which could reduce insulin’s neuroprotective effects despite systemic therapy. Interestingly, among the ten T1DM brains examined, only one was pathologically diagnosed with Braak stage 3-6 (excluded from this study). This low prevalence potentially supports insulin’s role in preventing p-Tau formation in AD. Indeed, a previous study has suggested that a combination of insulin and other diabetes medications may reduce p-Tau levels [4]. Whether insulin alone can prevent p-Tau formation and how it might be applied in this context remains an intriguing question in AD research.

Future experimental studies should investigate whether optimizing insulin delivery to the brain—such as through intranasal administration—can enhance its protective effects on cholinergic neurons and tau pathology. Additionally, exploring the interaction between peripheral and CNS insulin resistance in T1DM and their combined impact on neurodegeneration could provide critical insights. Such findings in animal models may guide the development of more effective therapeutic strategies targeting the underlying mechanisms of AD pathology.

Despite more frequent hypoglycemia in T1DM, both persons with T1DM and T2DM commonly experience hyperglycemia, which may explain our findings of lower GFAP-expressing astrocytes in both groups. Previous studies have also reported a loss of GFAP-positive astrocytes in other brain regions of T2DM subjects [18], although the mechanisms underlying hyperglycemia-induced astrocytic dysfunction require further investigation. Hyperglycemia’s adverse effects on blood vessels [16, 41, 51] and its association with increased alpha-SMA in AD [21, 56] were not observed in the NBM, suggesting that vascular dysfunction may not contribute to cholinergic neuronal loss in T1DM. Regarding other pathological changes associated with hyperglycemia, a previous study using streptozotocin-induced T1DM animal models found a lower production of ACh in striatal slices in culture, but did not detect difference in freshly isolated striata and hippocampal tissues [61].

While T2DM subjects showed comparable ChAT-ir to controls, epidemiological studies link T2DM with cognitive impairments [54]. Indeed, in subjects with Braak stage 3-6, we observed higher expression of CP13-ir, AT8-ir, and PHF-ir in T2DM subjects, indicating exacerbated neuropathological changes associated with AD progression. Elevated p-Tau levels have also been reported in the CSF of people with T2DM compared to healthy controls [34], consistent with our findings and epidemiological data. Moreover, our previous studies have observed less neurons that express proopiomelanocortin in the infundibular nuclei, fewer oxytocin neurons in the paraventricular nuclei (PVN), as well as reduced arginine vasopressin and vasoactive intestinal polypeptide in the SCN [8, 18, 26], suggesting that T2DM does affect neuronal function in various brain regions.

Microglia and the glymphatic system both function as keepers of brain homeostasis [7, 20, 23, 36, 59, 64], yet their roles in supporting cholinergic neuronal function remain unclear. Unlike activated microglia in AD brains [32], we observed a reduction in microglial cells in the NBM of people with T1DM. This reduction does not appear to be caused by hypoglycemia or hyperglycemia, since previous animal studies have shown that microglia become activated in response to hypoglycemia [63]. Moreover, in previous studies of our group, using diabetic *db*/*db* mice (known for uncontrolled hyperglycemia), we also did not observe a decrease in microglial cell density [12]. An alternative explanation could be that reduced cholinergic neuronal activity in the NBM leads to a decreased immune demand compared to normal physiological conditions, resulting in lower microglial activity over time and subsequently fewer microglial cells in the NBM. In contrast to microglia, studies investigating AQP4 levels in AD brains have produced conflicting results [46, 58]. Our finding of lower AQP4 expression in T1DM brains raises questions about whether reduced glymphatic activity is due to frequent hypoglycemia or decreased immune activity resulting from neuronal loss, as hypothesized for microglia alterations.

In conclusion, our finding of reduced ChAT-ir neurons in the NBM of people with T1DM, for the first time provides a potential mechanistic link between T1DM and AD. This finding also refers to the “cholinergic hypothesis,” one of the earliest theories of AD pathogenesis [3]. We suggest measuring cholinergic index, i.e., ratio of ChAT/AchE (acetylcholinesterase) in the CSF, as potential biomarker for diagnosing cognitive dysfunction in people with T1DM. If confirmed, acute testing with an acetylcholinesterase inhibitor may determine whether enhancing cholinergic neurotransmission improves cognitive performance. This may lead to considering the cholinergic index as a biomarker indicating early stages of cognitive decline. Additionally, enhancing cholinergic neurotransmission through supplementation with AChE inhibitors, such as Donepezil, which is used to treat symptoms of Alzheimer’s disease, could be considered as a therapeutic approach to improve cognitive performance in people with T1DM.

## Acknowledgements

We would like to thank Samantha E.C. Wolff, Rawien Balesar, Arja Sluiter, Joris Coppens and Roeland Lokhorst at the Netherlands Institute for Neuroscience, and Thuc-Anh Nguyen at the Tytgat Institute-Amsterdam UMC for their technical support.

## Author contributions

WJ, MJK and FCS performed morphological studies with human brain tissues. AK, DFS provided resources. AK, DFS, SES and CXY supervised the studies. The manuscript was originally written and edited by WJ, DFS, SES and CXY. All authors reviewed and edited the manuscript and had final responsibility for the decision to submit for publication.

## Consent statement

All authors have read and provided consent to be associated with this manuscript.

## Supplementary material

**Supplementary Table 1.** Antibodies information.

| Primary antibody | Source | Host | Catalog number | Specificity (PMID) | Dilution |
| --- | --- | --- | --- | --- | --- |
| Iba1 | Synaptic Systems | Rabbit | 234003 | 32814716 | 1:400 |
| GA130 | Netherlands Institute for Brain Research | Rabbit | α-HG-130 | 11222994 | 1:4000 |
| GFAP | DAKO | Rabbit | Z0334 | 32422642 | 1:1000 |
| AQP4 | Atlas antibodies | Rabbit | HPA014784 | 27893874 | 1:1000 |
| Alpha-SMA | Sigma Aldrich | Mouse | A5228 | 24024123 | 1:1000 |
| ChAT | Sigma Aldrich | Goat | AB1440P | 26792551 | 1:500 |
| AT8 | ThermoFisher | Mouse | MN1020 | 26792551 | 1:400 |
| CP13 | Gift from Fred van Leeuwen | Mouse | Peter Davies (1) | 24788298 | 1:400 |
| PHF-1 | Gift from Fred van Leeuwen | Mouse | Peter Davies (1) | 24788298 | 1:400 |

(1): Petry FR, Pelletier J, Bretteville A, Morin F, Calon F, Hebert SS, et al. Specificity of anti-tau antibodies when analyzing mice models of Alzheimer's disease: problems and solutions. PLoS One. 2014;9(5):e9425

## Supplementary Information for Methods

### Nissl staining

Sections were mounted on glass slides (superfrost+, Thermo Scientific) and dried on a 37°C heating plate. After 48 hours, the sections were deparaffinized in 100% xylene, rehydrated in grading ethanol (100% - 50%) and rinsed in distilled water. Next, the sections were submerged in a 0.5% thionine solution for 5 minutes and rinsed in water. After dehydration in graded ethanol (50%-100%) and xylene, sections were cover slipped using Entellan (Sigma-Aldrich, 107960, dried by air and ready for analysis.

### Heat-induced epitope retrieval

Sections were submerged in citrate buffer (pH 6.0) for the ChAT, Iba1, GFAP, alpha-SMA and citrate buffer (pH 9.0) for GA130. Afterward, sections were heated for 10 minutes by microwave treatment (700 W), and were subsequently cooled down for 30 minutes; for the CP13 and PHF1 antibodies, we performed epitope retrieval in room temperature by submerging the sections in formic acid (pH 2.0) for 10 minutes.

### Immunohistochemistry

After overnight primary antibody incubation, sections were rinsed in TBS and incubated for 60 minutes with biotinylated secondary antibody (1:400, Vector Laboratories). Then incubated for 60 minutes with avidin-biotin complex (1:800, Vectastain Elite ABC kit; Vector Laboratories Inc.) and were subsequently rinsed in TBS. Finally, sections were incubated in 0.5mg/ml 3,3’-Diaminobenzidine (Sigma Chemical Co., St. Louis, MO, DAB) in TBS. For ChAT and GA staining, 0.2% ammonium nickel sulphate and 0.01% H2O2 (Merck, Darmstadt, Germany) were added (DAB/Ni) (BDH; Brunschwig, Amsterdam, The Netherlands).

## Supplementary Figures and Legends

**Supplementary Fig. 1:**
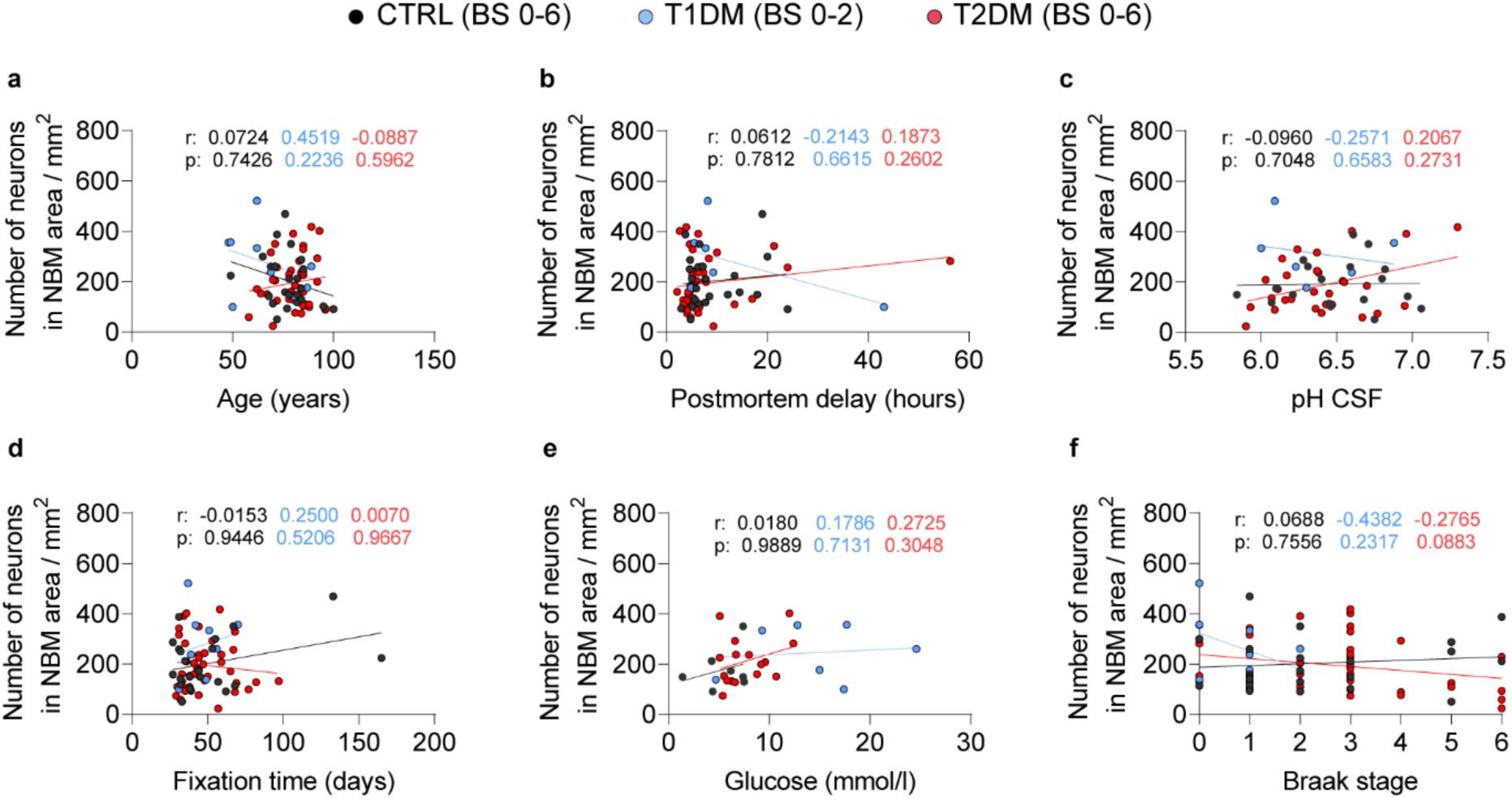
**a - f** Confounder analysis of the number of thionine-stained neurons in the in the nucleus basalis of Meynert (NBM) of control (CTRL), type 1 diabetes mellitus (T1DM) and T2DM subjects with Braak stage 0-2 or 0-6. No significant correlation was found between thionine-stained cells with these potential confounders.

**Supplementary Fig. 2:**
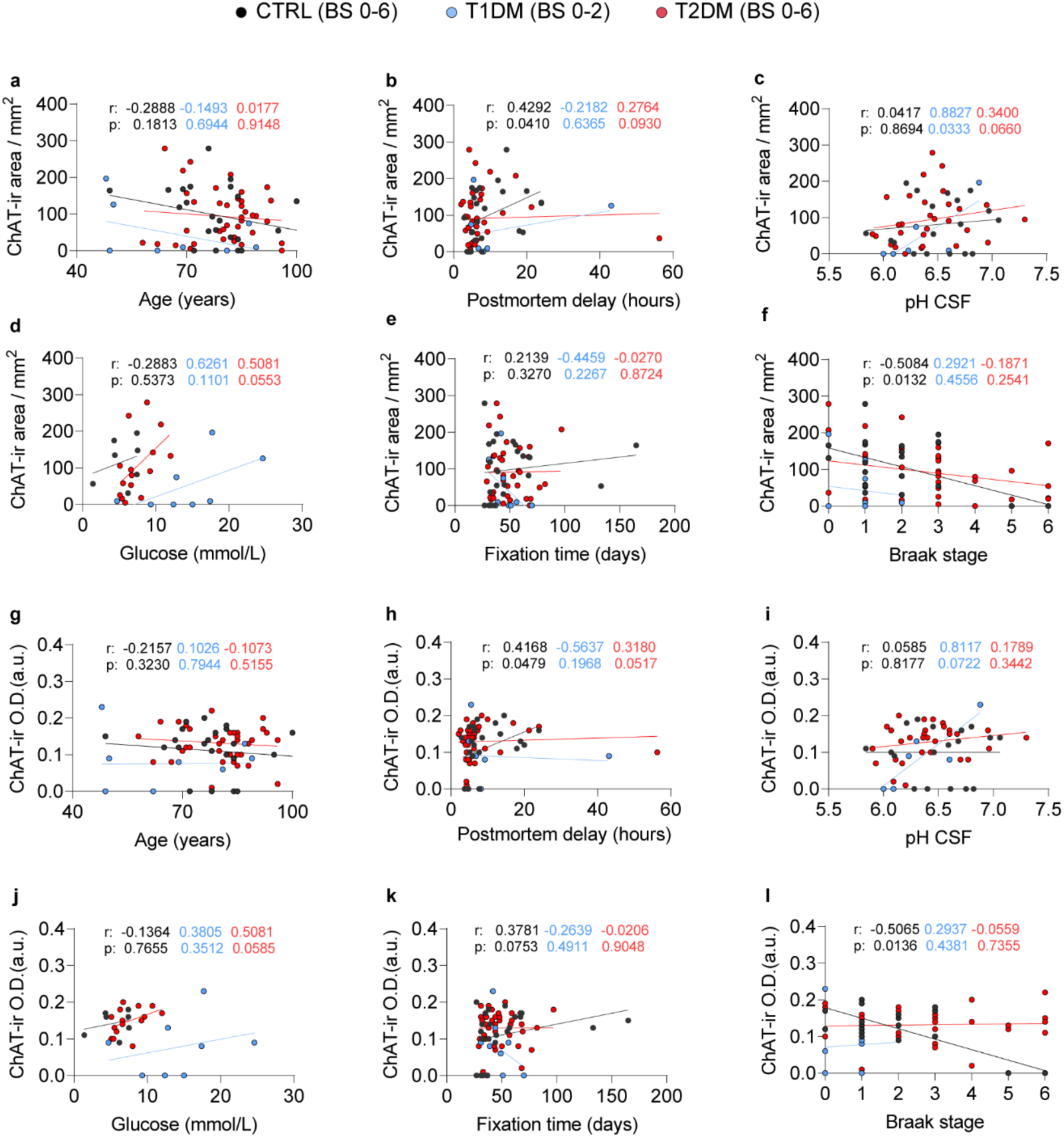
**a - l** Confounder analysis of choline acetyltransferase immunoreactive (ChAT-ir) areas and optical density (in arbitrary unit, O.D. (a.u.)) in the nucleus basalis of Meynert (NBM) of control (CTRL), type 1 diabetes mellitus (T1DM) and T2DM subjects with Braak stage 0-2 or 0-6.

**Supplementary Fig. 3.**
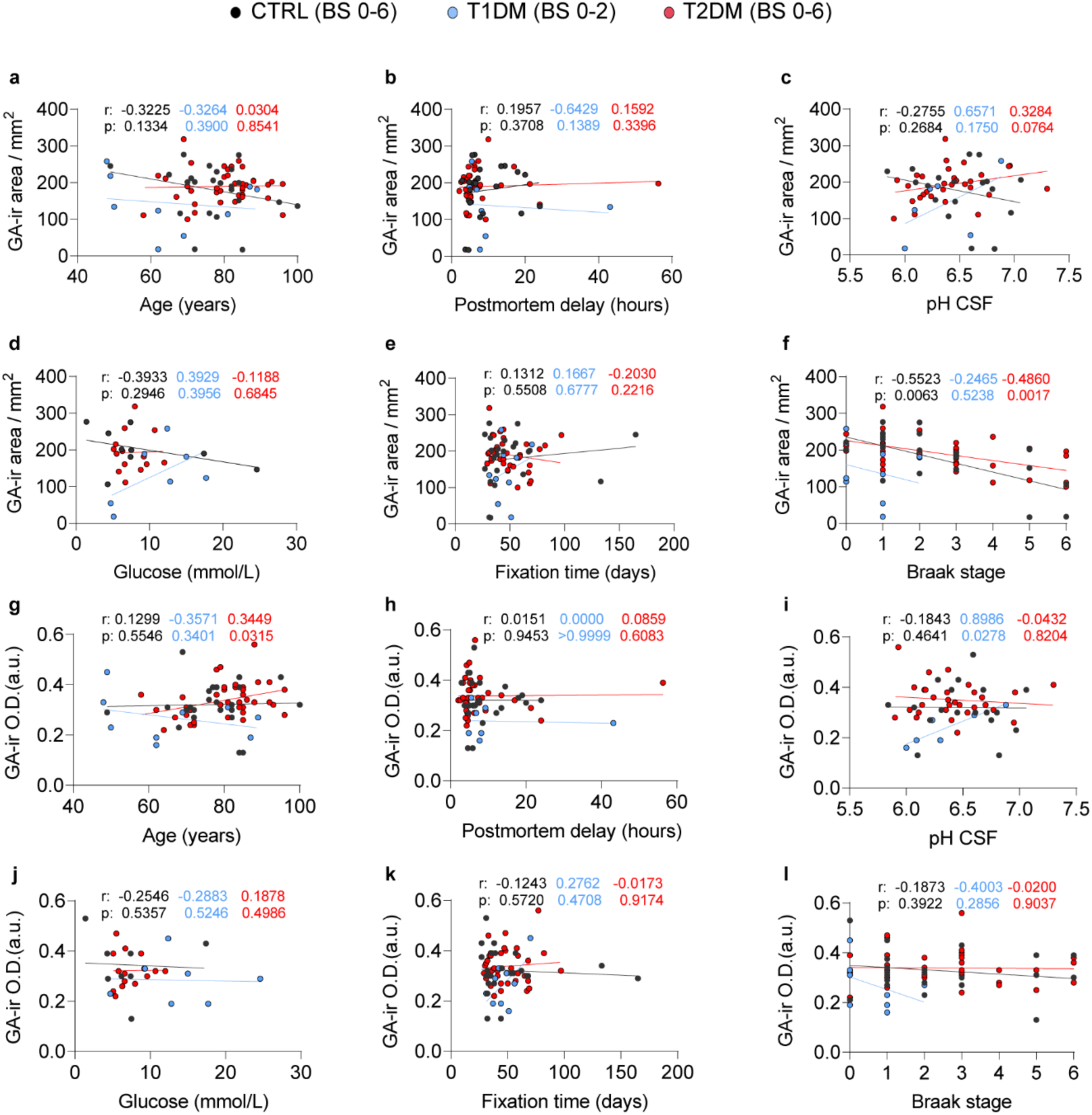
**a - l** Confounder analysis of Golgi matrix protein GA130 immunoreactive (GA-ir) areas and optical density (in arbitrary unit, O.D. (a.u.)) in the nucleus basalis of Meynert (NBM) of control (CTRL), type 1 diabetes mellitus (T1DM) and T2DM subjects with Braak stage 0-2 or 0-6.

**Supplementary Fig. 4.**
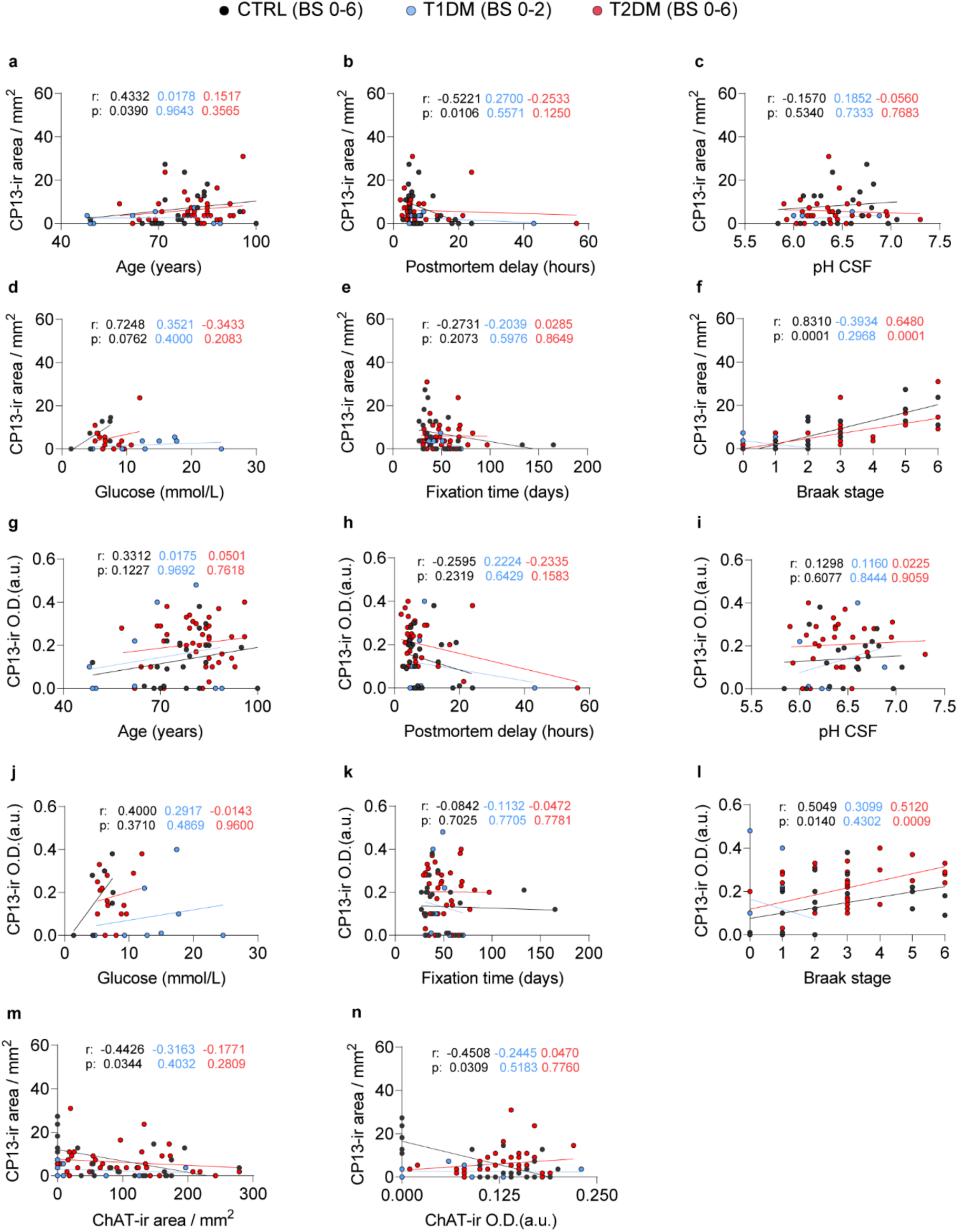
**a - l** Confounder analysis of p-Tau CP13 immunoreactive (CP13-ir) areas and optical density (in arbitrary unit, O.D. (a.u.)) in the nucleus basalis of Meynert (NBM) of control (CTRL), type 1 diabetes mellitus (T1DM) and T2DM subjects with Braak stage 0-2 or 0-6. **m, n** Correlation between the CP13-ir areas and the choline acetyltransferase immunoreactive (ChAT-ir) areas or O.D.

**Supplementary Fig. 5.**
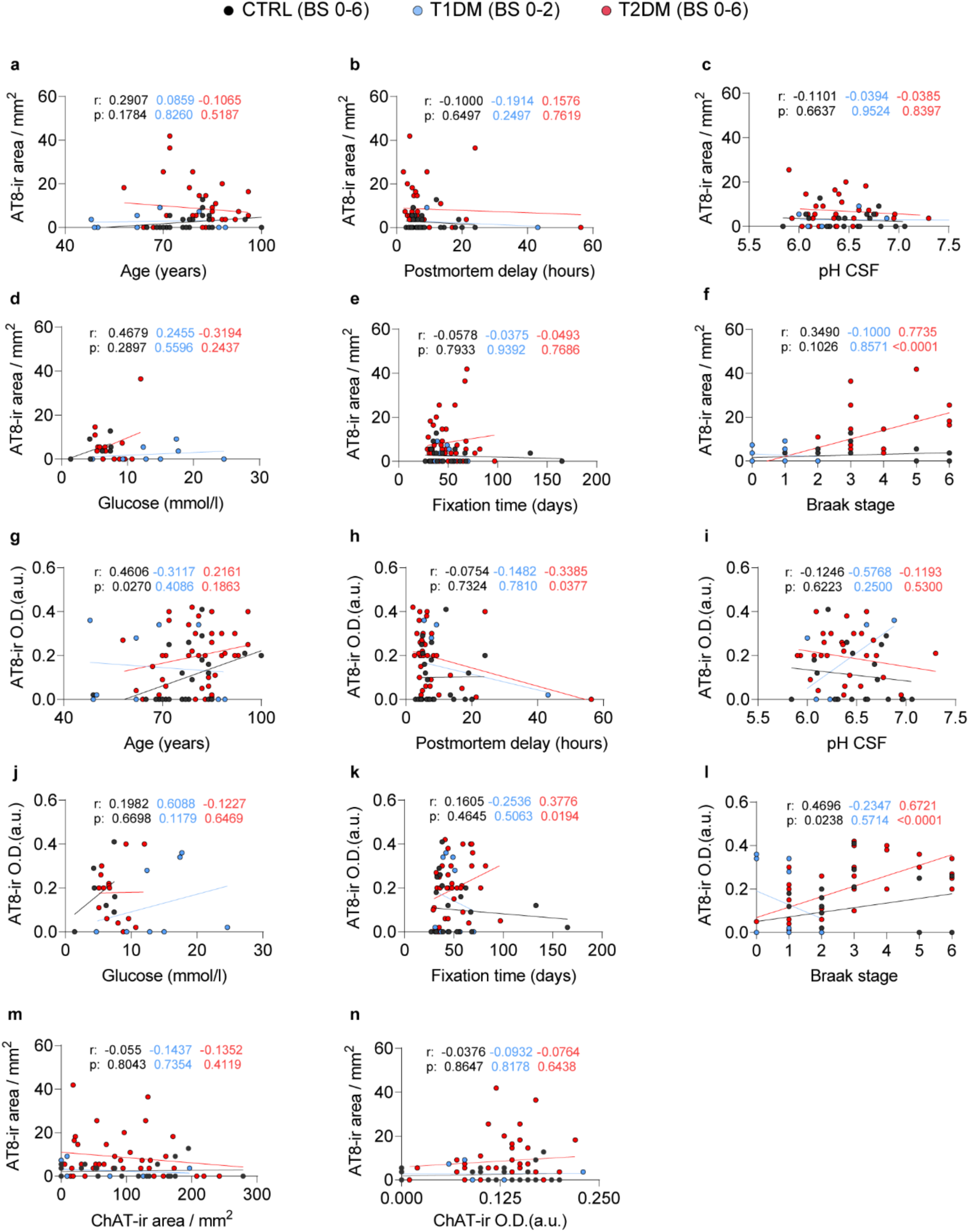
**a - l** Confounder analysis of p-Tau AT8 immunoreactive (AT8-ir) areas and optical density (in arbitrary unit, O.D. (a.u.)) in the nucleus basalis of Meynert (NBM) of control (CTRL), type 1 diabetes mellitus (T1DM) and T2DM subjects with Braak stage 0-2 or 0-6. **m, n** Correlation between the AT8-ir areas and the choline acetyltransferase immunoreactive (ChAT-ir) areas or O.D.

**Supplementary Fig. 6.**
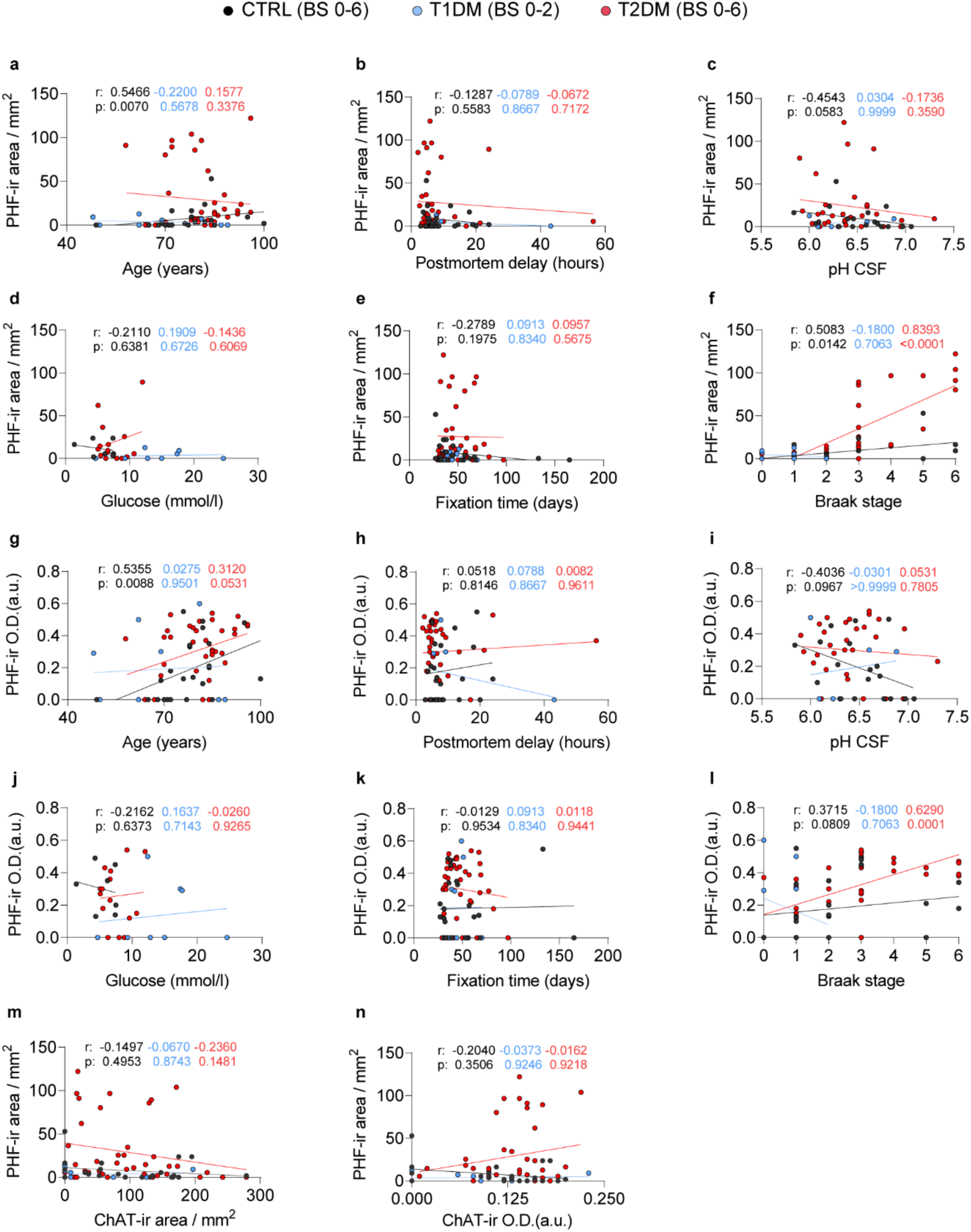
**a - l** Confounder analysis of p-Tau PHF immunoreactive (PHF-ir) areas and optical density (in arbitrary unit, O.D. (a.u.)) in the nucleus basalis of Meynert (NBM) of control (CTRL), type 1 diabetes mellitus (T1DM) and T2DM subjects with Braak stage 0-2 or 0-6. **m, n** Correlation between the PHF-ir areas and the choline acetyltransferase immunoreactive (ChAT-ir) areas or O.D.

**Supplementary Fig. 7.**
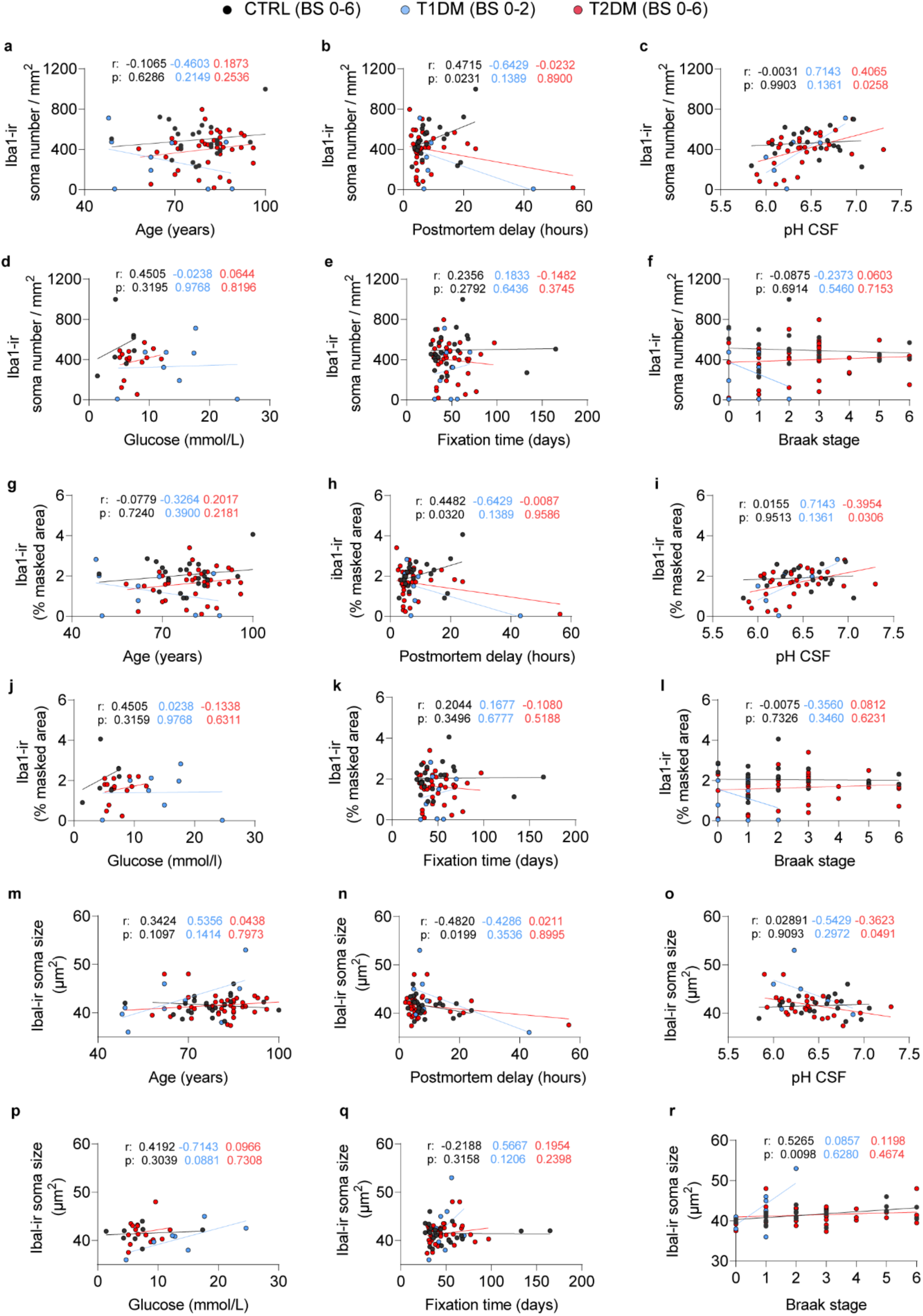
**a - r** Confounder analysis of ionized calcium binding adaptor molecule 1 immunoreactive (Iba1-ir) soma density, masked area (%) and soma size in the nucleus basalis of Meynert (NBM) of control (CTRL), type 1 diabetes mellitus (T1DM) and T2DM subjects with Braak stage 0-2 or 0-6.

**Supplementary Fig. 8.**
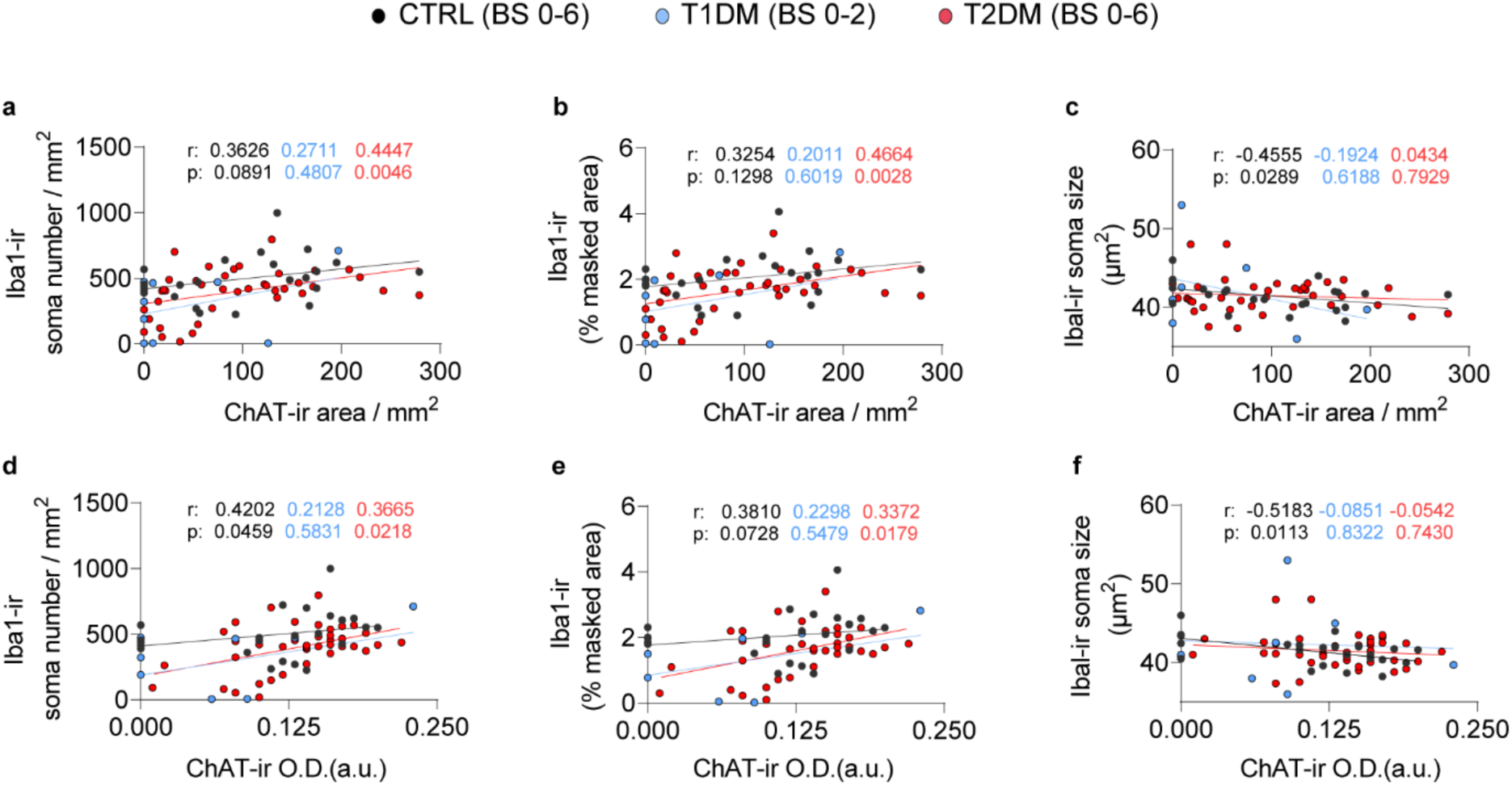
**a - f** Correlation analysis between the ionized calcium binding adaptor molecule 1 immunoreactive (Iba1-ir) soma density, masked area (%) and soma size with the ChAT-ir areas and optical density (in arbitrary unit, O.D. (a.u.)) in the nucleus basalis of Meynert (NBM) of control (CTRL), type 1 diabetes mellitus (T1DM) and T2DM subjects.

**Supplementary Fig. 9.**
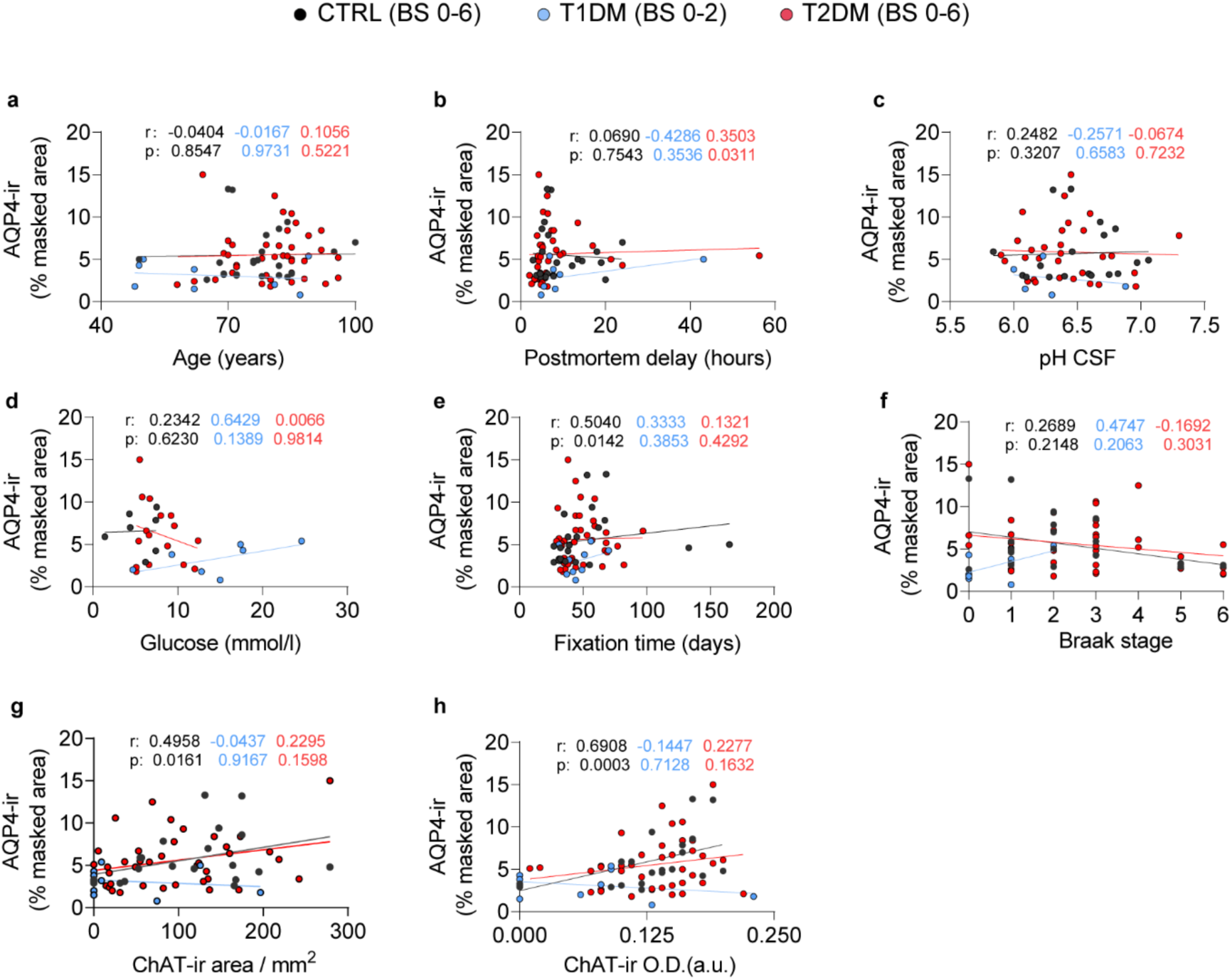
**a - f** Confounder analysis of aquaporin 4 immunoreactive (AQP4-ir) masked area (%) in the nucleus basalis of Meynert (NBM) of control (CTRL), type 1 diabetes mellitus (T1DM) and T2DM subjects with Braak stage (BS) 0-2 or 0-6. **g, h** Correlation between the AQP4-ir masked area (%) and the choline acetyltransferase immunoreactive (ChAT-ir) areas and optical density (in arbitrary unit, O.D. (a.u.)).

**Supplementary Fig. 1.**
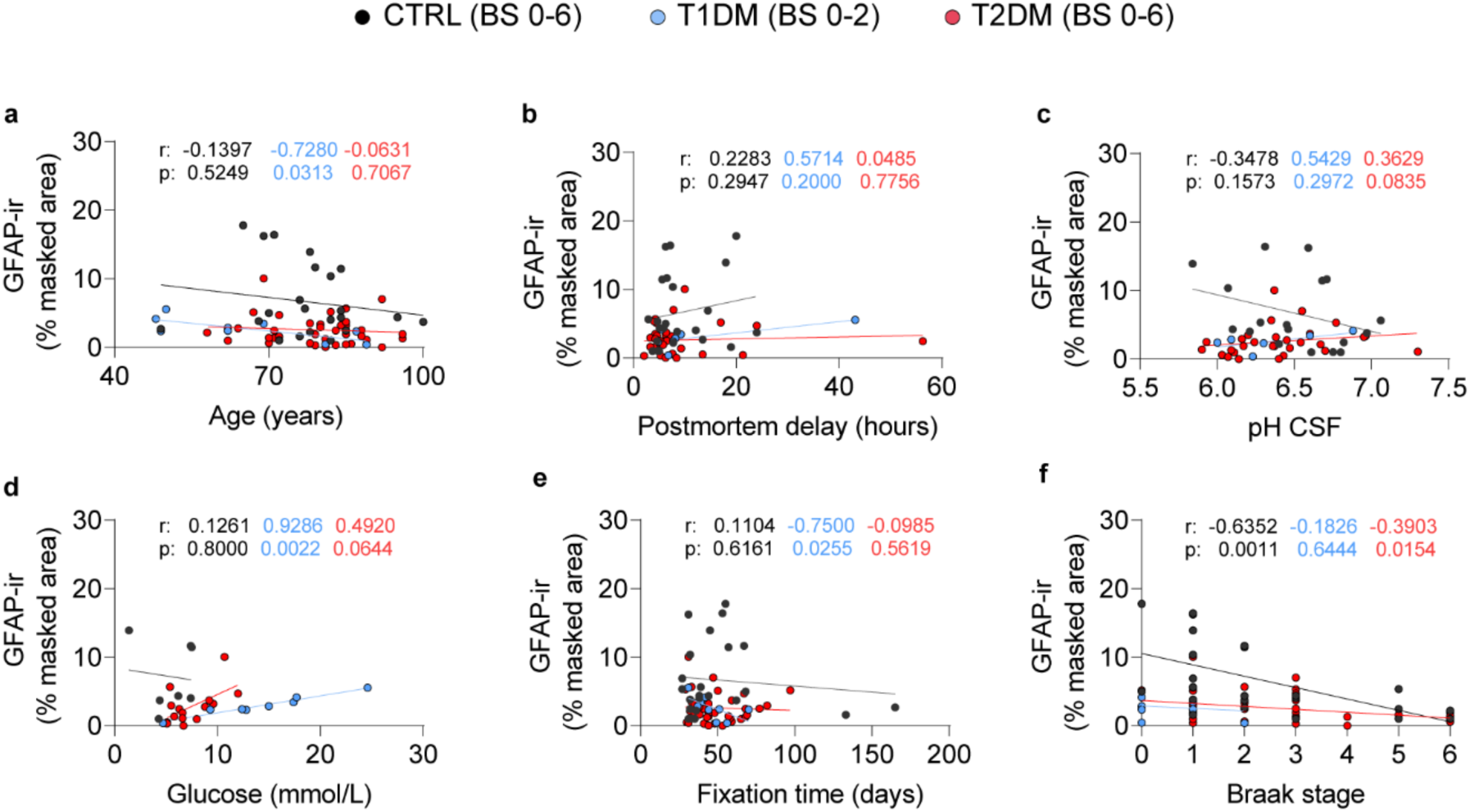
0 **a - f** Confounder analysis of glial fibrillary acidic protein immunoreactive (GFAP-ir) masked area (%) in the nucleus basalis of Meynert (NBM) of control (CTRL), type 1 diabetes mellitus (T1DM) and T2DM subjects with Braak stage 0-2 or 0-6.

**Supplementary Fig. 1.**
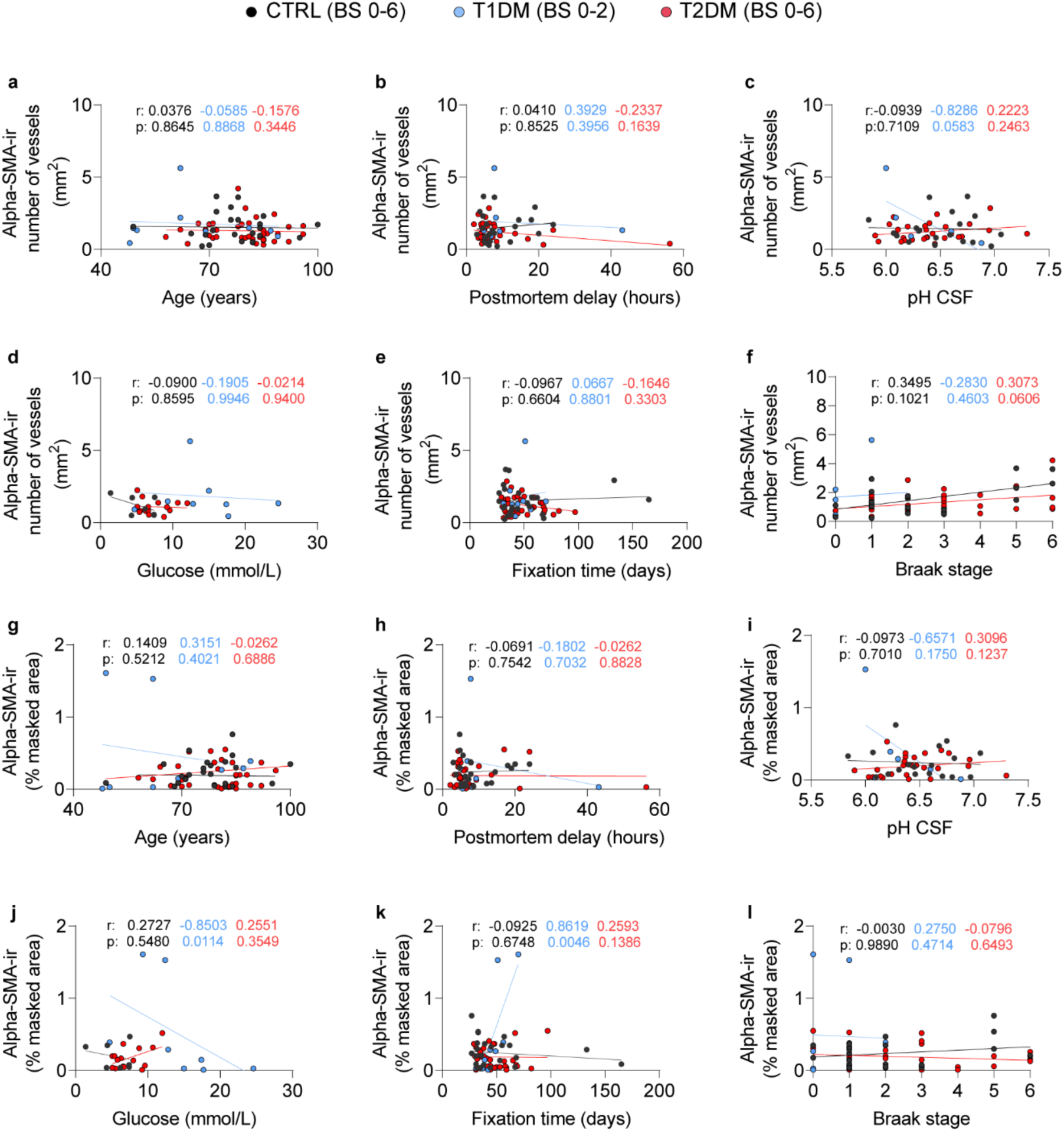
1 **a - l** Confounder analysis of alpha-smooth muscle actin immunoreactive (alpha-SMA-ir) vessel density and masked area (%) in the nucleus basalis of Meynert (NBM) of control (CTRL), type 1 diabetes mellitus (T1DM) and T2DM subjects with Braak stage 0-2 or 0-6.

